# Multi-modal, Label-free, Optical Mapping of Cellular Metabolic Function and Oxidative Stress in 3D Engineered Brain Tissue Models

**DOI:** 10.1101/2024.08.08.607216

**Authors:** Yang Zhang, Maria Savvidou, Volha Liaudanskaya, Varshini Ramanathan, Thi Bui, Matthew Lindley, Ash Sze, Ugochukwu O. Ugwu, Yuhang Fu, Matthew E. Dilsizian, Xinjie Chen, Sevara Nasritdinova, Aonkon Dey, Eric L. Miller, David L. Kaplan, Irene Georgakoudi

## Abstract

Brain metabolism is essential for the function of organisms. While established imaging methods provide valuable insights into brain metabolic function, they lack the resolution to capture important metabolic interactions and heterogeneity at the cellular level. Label-free, two-photon excited fluorescence imaging addresses this issue by enabling dynamic metabolic assessments at the single-cell level without manipulations. In this study, we demonstrate the impact of spectral imaging on the development of rigorous intensity and lifetime label-free imaging protocols to assess dynamically metabolic functions over time in 3D engineered brain tissue models comprised of human induced neural stem cells, astrocytes, and microglia. Specifically, we rely on multi-wavelength spectral imaging to identify the excitation/emission profiles of key cellular fluorophores within human brain cells, including NAD(P)H, LipDH, FAD, and lipofuscin. These enable the development of methods to mitigate lipofuscin’s overlap with NAD(P)H and flavin autofluorescence to extract reliable optical metabolic function metrics from images acquired at two excitation wavelengths over two emission bands. We present fluorescence intensity and lifetime metrics reporting on redox state, mitochondrial fragmentation, and NAD(P)H binding status in neuronal monoculture and the triculture systems to highlight the functional impact of metabolic interactions between different cell types. Our findings reveal significant metabolic differences between neurons and glial cells, shedding light on metabolic pathway utilization, including the glutathione pathway, OXPHOS, glycolysis, and fatty acid oxidation. Collectively, our studies establish a label-free, non-destructive approach to assess the metabolic function and interactions among different brain cell types relying on endogenous fluorescence and illustrate the complementary nature of the information that is gained by combining intensity and lifetime-based images. Such methods can improve understanding of physiological brain function and dysfunction that occurs at the onset of cancers, traumatic injuries and neurodegenerative diseases.

## Introduction

The coupling between energy metabolism and neurotransmission in the brain is critical for maintaining function at the organismal level. This is achieved, at least in part, by complex metabolic interactions between different brain cell types, leading to a highly heterogenous metabolic landscape over both space and time [1]. Tools that characterize the unique metabolic profiles of neurons, astrocytes, oligodendrocytes, and microglial cells and their interactions, are essential for understanding physiological brain function and disease mechanisms. Various non-invasive methods, such as PET, fMRI, NMR spectroscopy [2–4], and optical imaging [5], provide important insights into human brain metabolism and connectivity, but lack the resolution to capture single-cell metabolic diversity. Single-cell transcriptomic and mass-spectrometry-based metabolomic studies offer detailed metabolic profiling at the cellular level [6, 7]. However, these powerful methods are destructive, require extensive processing, and cannot capture dynamic changes. Thus, methods that can yield metabolic insights non-destructively and with single cell resolution could significantly impact our understanding of the heterogeneous and dynamic functions and interactions of brain cells.

High-resolution two photon imaging of living animals has provided unique insights into various aspects of brain function, including spontaneous behavior [8], memory formation and learning [9], and inflammatory responses of specific cell types, such as astrocytes and microglia [10]. Most of these studies have relied on exogenous fluorescent labels, which are sensitive to Ca^++^ to assess neuronal or astrocytic firing or are stably expressed in certain cell types. Beyond Ca^++^ sensitive probes, a wide range of genetically encoded fluorescent indicators (GEFIs) that serve as sensors of metabolic cofactors (NAD+, NADH), sugars (glucose/galactose), monocarboxylates (pyruvate, lactate), amino acids (arginine, glycine, serine), and energy nucleotides (ATP, ADP) have been developed and used to perform dynamic metabolic assessments with single cell resolution [11]. However, GEFIs rely on viral vector manipulations, which have limited success with primary human cells, require careful calibrations, and provide access to a small fraction of the metabolome. Label-free, two photon excited fluorescence imaging of brain metabolism has also been reported in brain slice preparations and in vivo in animals providing a unique opportunity to study dynamic metabolic changes with single cell resolution without any requirements for cell manipulations [12–15].

Assessments of dynamic brain metabolic function changes relying on the endogenous fluorescence of NAD(P)H and flavins were among the seminal studies performed by Britton Chance and his colleagues several decades ago, which characterized both the natural fluorescence of these co-enzymes and established their key roles in several metabolic pathways [16–20]. We use “NAD(P)H” to refer to fluorescence emanating from both NADH and NADPH, since their fluorescence intensity excitation/emission characteristics are indistinguishable. We refer collectively to signals emanating from FAD and flavoproteins, such as LipDH, considered the major contributor to the flavin signals detected from cells and tissues, as flavoprotein (FP) signals [21]. The redox ratio, defined in our studies, as the ratio of FP/(NAD(P)H+FP) fluorescence, is used as a metric of the oxidoreductive state of the cells, consistent with the definition established by Chance [22]. We note that the optical redox ratio is defined in different ways in our field (e.g., NAD(P)H/FP, NAD(P)H/(FP+NAD(P)H)). We use FP/(NAD(P)H+FP) because the values are lie between 0 and 1 enabling unambiguous comparisons of data acquired over multiple time-points. In addition, this redox ratio definition is consistent with the notion of redox potential, often used in biochemistry, so that a higher redox ratio is typically associated with higher levels of oxidative phosphorylation (OXPHOS) than glycolysis [23–25]. In addition to the redox ratio, other metrics of metabolic function can be derived from label-free two photon images. Specifically, we have shown that the intensity fluctuations in NAD(P)H images, which are typically acquired with high numerical aperture (NA) objectives for two-photon imaging, can be analyzed to provide a metric of the level of mitochondrial fragmentation or clustering in a wide range of specimens, including living humans [23, 26–30]. This approach relies on the fact that the quantum efficiency of NAD(P)H fluorescence is enhanced two to ten-fold when NAD(P)H is in its bound vs its free form, with bound NAD(P)H being much more prevalent in the mitochondria rather than the cytosol. This approach allows us to assess dynamic changes in mitochondrial fusion and fission as a complementary metric of metabolic function [31]. Finally, fluorescence lifetime imaging, especially of NAD(P)H, is highly sensitive to metabolic function as it reflects changes in binding state, and other microenvironmental factors, such as pH and viscosity [32, 33]. Typically, enhanced levels of NAD(P)H in the bound state, corresponding to increased contributions from long-lifetime fluorescence components, are associated with enhanced levels of OXPHOS [23]. Despite their high sensitivity to alterations in metabolic function, these metrics lack specificity, which is a limitation of label-free, optical metabolic imaging. However, the combined consideration of all these parameters may provide some specificity to the interpretation of the origins of the observed optical changes [23, 34].

In this study, we demonstrate how label-free spectral imaging can improve the analysis of multi-modal (intensity and lifetime) TPEF images to extract in a robust manner brain cell metabolic function insights. Our experiments are performed with a set of novel in vitro engineered brain tissue models, comprising a silk-collagen hydrogel seeded with human induced neural stem cells (hiNSCs), astrocytes and microglia [35, 36]. While these tissues have obvious limitations in terms of representing the complexity of fully functioning brains, they include human cells and can be used to address specific questions in a more highly controlled environment that is more physiologically relevant than 2D cultures [37, 38]. The silk-hydrogel-based tissue models we use have been shown to mimic important aspects of neuroinflammation following traumatic brain injury [37–39], plaque formation associated with Alzheimer’s disease [40–42], and physiologically relevant responses to various treatments. Here, we show that spectral images can be analyzed to establish the detailed excitation/emission profiles of four key cellular fluorophores, NAD(P)H, LipDH, FAD and lipofuscin. Lipofuscin is a complex mixture of highly oxidized cross-linked macromolecules, such as proteins, lipids, and sugars, and has been shown to yield bright autofluorescence (relative to other endogenous fluorophores), with broad and variable excitation/emission characteristics [43]. It is usually the result of interactions with reactive oxygen species that are naturally produced in the cell, especially during OXPHOS. As a result of the overall high need for efficient energy production [44, 45], lipofuscin levels in brain cells are high. Ignoring its contributions to the emission within excitation/emission wavelength ranges that we attribute to NAD(P)H and flavins has significant implications in terms of the rigor with which the optical metrics we derive can report cellular metabolic function. Knowledge of the full spectral characteristics of the fluorophores enables the development of algorithms to identify lipofuscin pixels so that they can be removed from further analysis to extract signals that can be reliably attributed to NAD(P)H and flavins. Our studies highlight the presence of significant metabolic function differences between neurons and glial cells (astrocytes and microglia). They provide evidence for the high proportion of NADPH relative to NADH in these cells and illustrate how more specific metabolic insights can be gained by combining metrics associated with redox state, mitochondrial fragmentation, and NAD(P)H binding status.

## Results

### Approach for long-term metabolic profiling of cell interactions in engineered brain tissue models

To establish a robust analysis platform to assess metabolic function and interactions among different types of human brain cells from label-free two photon images, we utilize a novel three-dimensional engineered brain tissue model, based on a silk-collagen hydrogel embedded either with just human induced neural stem cells (hiNSCs), which we call the *monoculture* model (Mono), or with hiNSCs, primary human astrocytes, and HMC3 human microglial cells, referred to as the *tri-culture* or *NAM* tissue model(Fig. 1A) [35, 36]. The high spatial resolution and non-destructive nature of the two-photon imaging approaches enable comparisons of results from the monoculture and NAM tissue models, highlighting their importance for assessing metabolic cell interactions. Additionally, these assessments underscore the physiological relevance of the engineered brain tissue models and demonstrate their potential for enabling studies that are not feasible with human tissues. To account for sample heterogeneity, we select three distinct imaging regions from each scaffold (center, middle, and edge; Fig. 1Ac) and perform two-photon imaging on the same batch of samples weekly or bi-weekly for up to three months (Fig.1Ad). In the first set of experiments, we employ spectral two-photon excited fluorescence (TPEF) imaging at 755 and 860 nm excitation, to identify the key endogenous fluorophores and their corresponding spectral excitation/emission profiles through spectral decomposition (Fig. 1B). One of these fluorophores, lipofuscin, is known to have intense, broad fluorescence and can confound the accuracy of measurements focused on emission from NAD(P)H and FPs, which are typically used to assess metabolic activity. Availability of the chromophore spectra enables the development of a robust protocol to analyze label-free TPEF images acquired at 755 and 860 nm at two emission bands (500-550 nm and 435-485 nm, represented by 525 and 460 in Fig. 1Cb-c, f-g) to quantify lipofuscin, NAD(P)H and FPs. For this set of experiments, we manually annotate cell types (Fig. 1Ci) using a combination of morphological and functional characteristics. To distinguish microglia from astrocytes during annotation, HMC3 cells are labeled with mCherry, emitting fluorescence collected by a 604–644 nm detector, indicated by “Red” (Fig. 1Cd). This labeling serves as a definitive microglial cell marker and does not interfere with our two emission bands. We rely on a ratiometric approach to extract the optical redox ratio, defined as FP/(NAD(P)H + FP) [23], using the corresponding NAD(P)H and FP signals (Fig. 1De), and a Fourier-based analysis approach of the TPEF NAD(P)H intensity fluctuations to quantify mitochondrial fragmentation levels (Fig. 1Dd). Finally, we acquire fluorescence lifetime images and employ phasor analysis to examine in detail NAD(P)H fluorescence lifetime characteristics [46]. Thus, combined with the TPEF intensity-based metrics, we gain insights into the differential utilization of different metabolic pathways by neurons and glial cells, including NADPH utilization, OXPHOS, glycolysis, and fatty acid oxidation (Fig. 1E).

**Fig 1.**
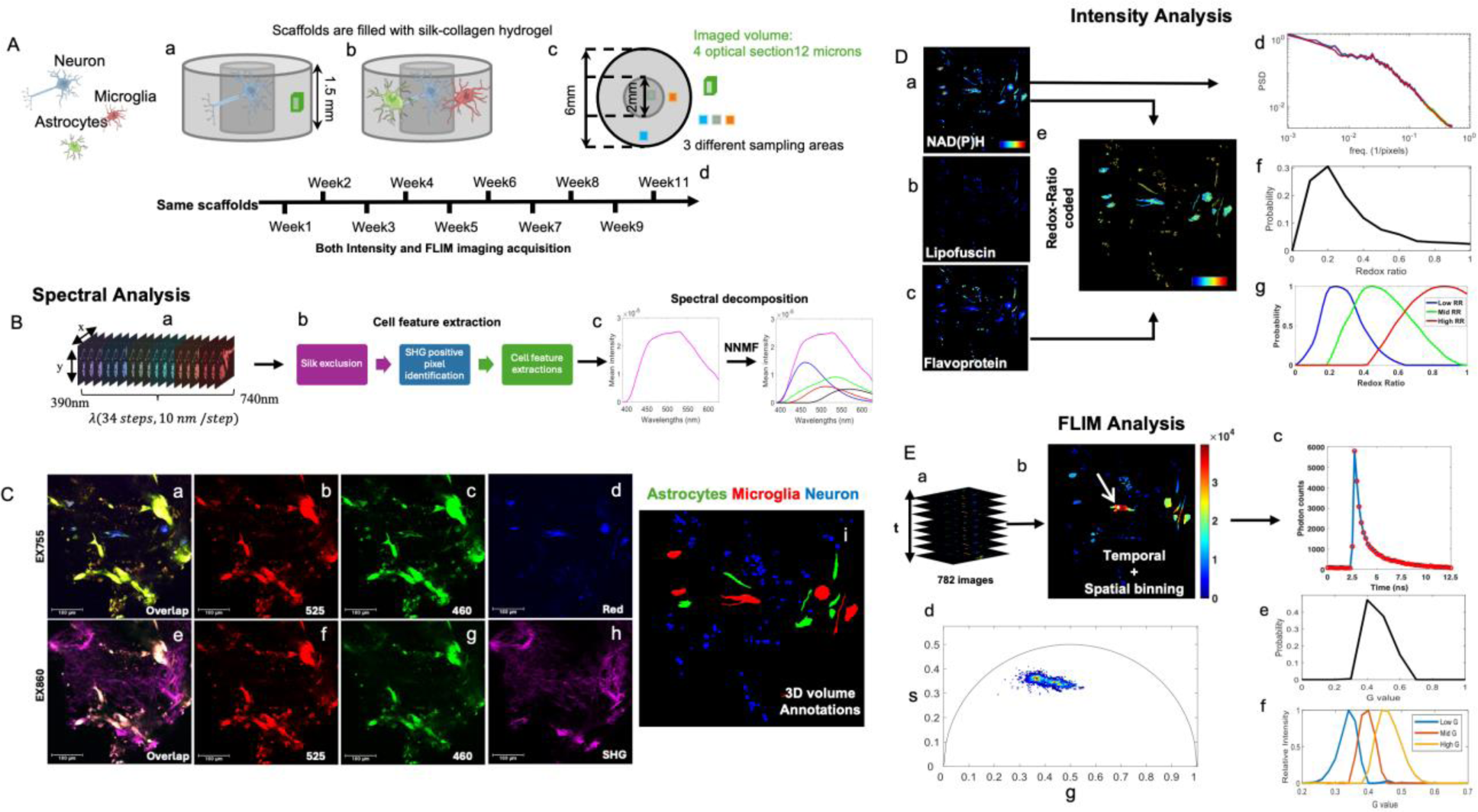
Experimental setup (A) and analytical approach (B-E)). (A) Schematic representation of 3D neuronal monocultures (Aa) and tricultures (Ab), along with imaging areas and sampling volumes (Ac) indicated by green, blue, and orange squares to capture heterogeneity. Ad shows imaging acquisition periods for both tissue types. (B) Spectral data acquisition involves 755 nm and 860 nm excitation, covering 390 nm to 740 nm emission (Ba), with a workflow for cell feature extraction from spectral data (Bb) and spectra decomposition using non-negative factorization to identify key fluorophores (Bc). (C) Excitation/emission channels include images acquired at 755 nm (Ca-d) and 860 nm (Ce-h), with overlapped images from 755 nm (Ca) and 860 nm (Ce), and manual annotations for cell features (Ci). (D) Intensity-based analyses feature NAD(P)H images at 755ex/460em (Da), lipofuscin images at 860ex/460em (Db), flavoprotein images at 860ex/525em (Dc), redox ratio (RR) color-coded images (RR=FP/(NAD(P)H+FP)) generated from Da and Dc (De), mitochondrial clustering analyses from Da (Dd), redox ratio distributions from redox ratio maps (Df), and decomposed redox distribution components (Dg). (E) FLIM analyses include fluorescence lifetime data illustration (Ea), temporal and spatial binning (Eb) with a color bar representing photon counts, a lifetime decay trace from a single pixel highlighted in white and pointed by the white arrow in Eb. (Ec), phasor analysis (Ed), g distribution derived from g values in Ed (Ee), and g component evaluation (Ef). All images in C-E are maximum projections of a 3D volume from the same ROI. Ca-h images are pseudo-colored; Da-d images use the ‘jet’ colormap, where blue represents lower values and red represents higher values. The color scale for Da-c ranges from 0 to 0.001, while Dd ranges from 0 to 1.

### Decomposition of spectral images identifies key fluorophores and distinct excitation/emission profiles

We use two excitation wavelengths, 755 nm, and 860 nm, to capture contributions from NAD(P)H and FPs, essential for metabolic function assessments. However, the porous silk hydrogel and the collagen fibers also exhibit strong autofluorescence due to cross-links (Fig.1C) [47]. We rely on the strong autofluorescence of silk scaffold features to manually exclude such contributions and define the region of interest (ROI) where cells reside, utilizing the maximum projection for each imaged volume (Fig. S1E). Then, we use an Image Segmentation Baseline Algorithm (ISBA) to identify cells (supplemental methods). Briefly, we focus on the second harmonic generation (SHG) images acquired at the 860 nm excitation/430 nm emission to identify intense SHG pixels, typically associated with strong collagen cross-link autofluorescence. We extract cell features separately in regions with strong and weak SHG containing pixels (Supplemental methods, Fig. S1-S3). Spectra in Fig. 2A-B are the peak normalized mean profiles of the cells imaged each week. For each ROI, the total cellular intensity is normalized to the cell pixels identified in that field. Red lines represent spectra from neuronal monocultures, while blue lines represent the mean spectra from the NAM system, where all three cell types are present. We observe significant differences between monoculture and NAM tissues under 755 nm excitation, which can be attributed to relative variations in NAD(P)H and FP content, both efficiently excited at this wavelength. For both cultures, we also note time-dependent red shifts. For 860 nm excitation, we do not expect measurable NAD(P)H excitation efficiency [48]. Thus, the time-dependent shifts at this wavelength suggest the presence of more than one fluorophore.

**Fig 2.**
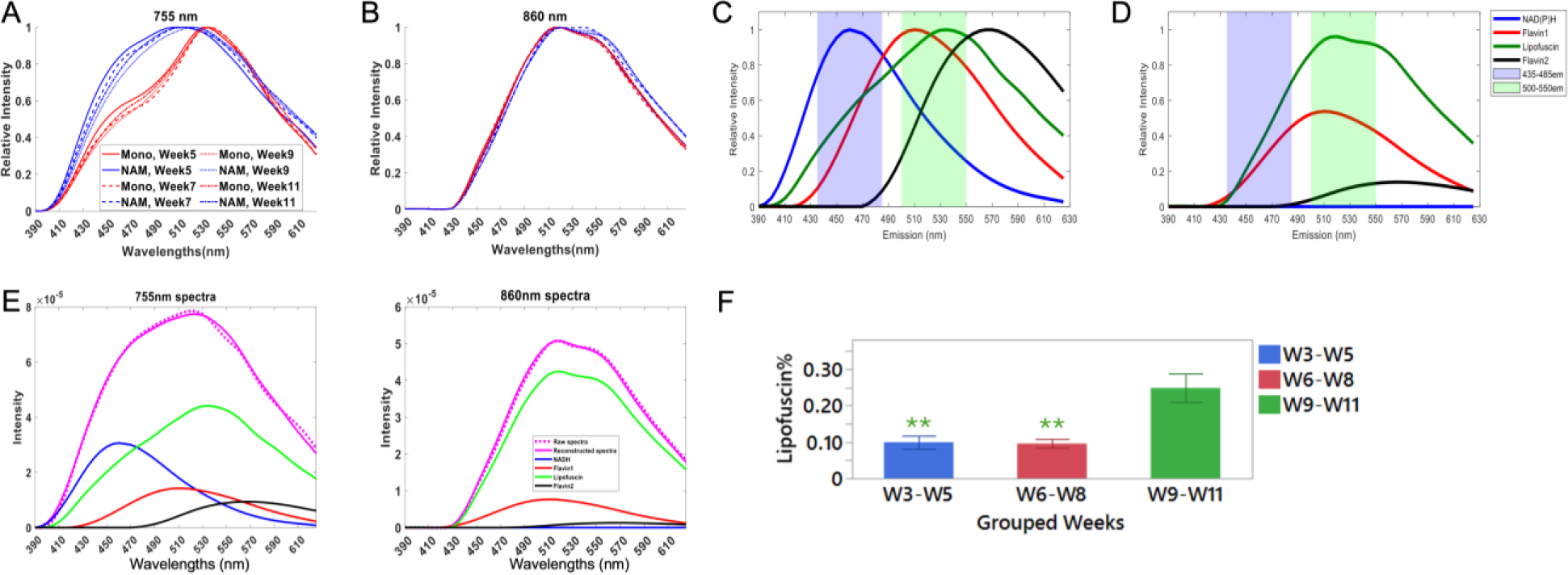
Identification of key human brain cell endogenous fluorophores and their excitation/emission characteristics. Cell spectra from neuronal monocultures (Mono-red) and tri-cultures (NAM-blue) acquired at 755nm (A) and 860nm (B) excitation wavelengths at 5,7,9 and 11 weeks from the onset of tissue culture. Spectrally decomposition cellular components at 755ex (C) and 860ex (D), attributed to NAD(P)H-blue, LipDH-red, FAD-black, and lipofuscin-green. (E). Representative NAM spectra from a sample 9 weeks in culture (solid magenta) with corresponding fit (dashed magenta), reconstructed using derived component shapes and their corresponding weights, showing the contributions from each fluorophore. Lipofuscin is a dominant contributor at both excitation wavelengths. (F) Spectrally deconvolved lipofuscin amounts from neuronal monocultures indicating a significant increase, especially during the last month of culture.

To identify the number of key fluorophores and their associated spectra, we use non-negative matrix factorization (NNMF) to analyze the combined spectra from 755 nm and 860 nm excitation obtained from the same locations. To facilitate meaningful spectral extraction, we rely on three assumptions: (a) negligible NAD(P)H TPEF excitation at 860 nm based on previous studies [48], (b) similar NAD(P)H spectral profiles in epithelial cells and brain cells since NAD(P)H participates in similar cellular reactions, and (c) consistent emission spectra of fluorophores such as NAD(P)H and FPs under different excitation wavelengths (i.e. 755 and 860 nm) (Supplemental Methods). This analysis indicates that excellent fits to the measured spectra can be acquired when we utilize a minimum of four components attributed to NAD(P)H, lipofuscin and two flavins (Fig. 2C-D, and Fig. S4-S6). The NAD(P)H component is derived from analysis of highly proliferative epithelial cells, rich in NAD(P)H [49]. The two FP shapes are determined based on their 860 nm profiles since NAD(P)H crosstalk is minimal at this excitation wavelength. Specific attribution to lipoamide dehydrogenase (LipDH) and flavin adenine dinucleotide (FAD) is based on established excitation/emission spectral and lifetime differences [48, 50] (Fig. S6). The extracted lipofuscin shape is characterized by broad emission and highly efficient excitation at both wavelengths. Since lipofuscin is considered a product of oxidative stress, we validate the identity of its spectral features with experiments that induce oxidative stress in microglial cultures (Fig. S7 and supplemental materials).

As shown in Fig. 2C-D, our analysis yields the wavelength-dependent excitation efficiency of each fluorophore. The combination of these four spectral components accurately describes the variations in all spectra detected from both the neuronal and NAM brain tissues (N = 174), with a residual of 0.03, meaning 96% of the variations are captured (Fig. 2E). Lipofuscin contributions are significant for both excitation wavelengths and they increase with culture time (Fig. 2F). Therefore, removing lipofuscin fluorescence from signals that we typically associate with NAD(P)H and flavins is important for accurate metabolic function assessments. While spectral emission features from the four components overlap, there are significant differences. Importantly, each component has very distinct excitation efficiencies at 860 nm versus 755 nm of ∼0:1 for NAD(P)H, ∼0.54:1 for LipDH (flavin 1), ∼0.14:1 for FAD (flavin2), and ∼0.96:1 for lipofuscin. These differences are exploited to establish robust fluorophores quantification approaches from images at two emission ranges (shaded areas in Fig. 2C-D) for each excitation wavelength.

### Spectra inform extraction of more accurate fluorophore contributions in TPEF images from a limited number of excitation/emission wavelengths

In most label-free metabolic imaging TPEF studies, a combination of two excitation and emission wavelength ranges are used to collect signals efficiently by non-descanned detectors that are typically attributed to NAD(P)H and flavins. In Fig. 2C-D, we overlay with the fluorophore spectra the shaded wavelength regions that correspond to the bandpass filters that we typically use to collect NAD(P)H (435-485nm emission, 755 nm excitation) and flavin (500-550 nm emission, 860 nm excitation) TPEF (Fig. 2C-D and Fig. S8). Wavelengths lower than 800 nm excitation are typically used to collect NAD(P)H, with the emission centered around the 450-460 nm peak and narrow enough to prevent significant contributions from flavin-associated TPEF, which is also efficiently excited at these wavelengths. Illumination longer than 850 nm is used to excite, more specifically, flavin TPEF, typically collected at wavelengths beyond 500 nm. Thus, given the spectra of the fluorophores and the emission ranges collected by the non-descanned detectors in our imaging system, we selected 0.8 as a relatively conservative threshold in the ratio of the intensities collected by the 500-550 nm detector at 860 and at 755 nm, to identify pixels with significant lipofuscin contributions. We note that in some cases, the overall intensity of those pixels for a given excitation/emission configuration may not be very high because the lipofuscin content is low.

Following lipofuscin pixel removal, we assume that the remaining signal is attributed to NAD(P)H and flavins. While NAD(P)H contributions are dominant in the 435-485 nm detector at 755 nm excitation, there are significant potential contributions by LipDH (flavin 1), especially for cells such as neurons with highly active OXPHOS, which is fueled by NADH consumption and can have similar NAD(P)H and flavin fluorescence levels. Given the full spectra we can accurately quantify the relative contributions of each fluorophore from the four images acquired in the two emission regimes at the two excitation wavelengths, as described in the “crosstalk adjustment section” of the supplemental materials (Fig. S8). We verify the accuracy by comparing the intensity-based and spectral-based redox ratio calculations from the same set of neuronal monocultures (Fig. S9). Finally, we validate the sensitivity of the NAD(P)H intensity, mitochondrial clustering and redox metrics as extracted following this approach with a set of hypoxia measurements performed using microglial cell cultures (Fig. S10) [23, 26].

### Endogenous TPEF intensity-based metabolic assessments reveal differential metabolic profiles and interactions between neurons and glial cells

Representative redox ratio-coded projections are shown from tricultures and monocultures following 3,7, and 11 weeks of culture. Neurons exhibit higher redox ratios (Fig. 3A-D) and lower mitochondrial fragmentation (Fig. 3E) than astrocytes and microglia, indicating greater OXPHOS and/or glutaminolysis activity [23]. The higher lipofuscin levels in neurons (Fig. 3F) indicate increased oxidative stress, likely because of their high reliance on OXPHOS for energy production, which generates ROS. Neurons also have limited anti-oxidant capabilities due to their lower GSH and NADPH pools compared to astrocytes [51] [1]. Although overall similar, there are some differences in the intensity-based metabolic function readouts extracted for astrocytes and microglia. For example, during weeks 6-8, the lower mitochondrial fragmentation level in astrocytes compared to microglia suggests increased glutaminolysis and/or OXPHOS in astrocytes (Fig. 3E). Interestingly, astrocytes but not microglia show an increase in lipofuscin levels at weeks 9-11 (Fig. 3F), indicating greater oxidative stress, possibly because of elevated OXPHOS activity during the earlier time point. In principle, we expect microglia to utilize OXPHOS as a primary means to meet energy demands [52]. However, in these tissues, microglia appear metabolically more similar to astrocytes than to neurons. This could indicate some functional limitations for the type of microglia used in this model, which have been reported previously [53]. Other types of human microglia, such as induced pluripotent stem cell derived microglia, may be used to improve relevance to tissue function [35].

**Fig 3.**
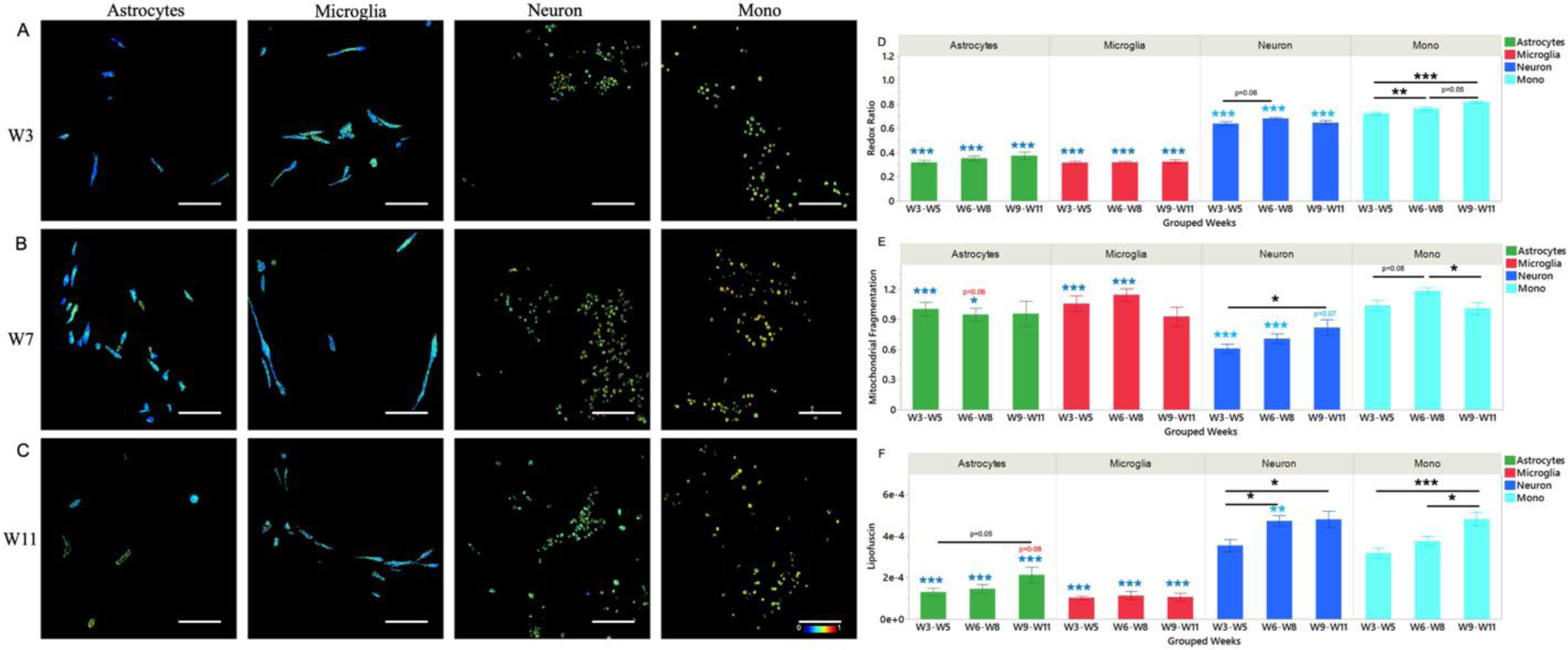
Intensity-based optical metabolic readouts. (A-C) Maximum projections of representative redox ratio color-coded images for different cell populations at Week 3 (A), Week 7 (B), and Week 11 (C). All images share the same color bar and scale bar. (D) redox ratio of different cell types as a function of time. Weeks were separated into three groups: W3-W5 (ROIs acquired at Weeks 3, 4, and 5), W6-W8 (Weeks 6, 7, and 8), and W9-W11 (Weeks 9 and 11). (E) Mitochondrial clustering comparisons. (F) Lipofuscin intensity comparisons. (*, **, and *** denote p<0.05, p<0.01, and p<0.001, respectively). Weekly data are categorized into three ranges, each cell type represented by a distinct color. bar. (black asterisks: differences for each cell group among the three time points, and color asterisks: differences for comparisons at a given time point among Neurons, astrocytes, microglia (ANOVA), and between Neurons in Mono and Neurons in NAM)

In monocultures, neurons show higher redox ratios relative to neurons in NAM, but overall levels of fragmentation are also higher, suggesting that mitochondria are highly active, but may be becoming stressed. Interestingly, NAM neurons exhibit very limited changes in redox ratio from W3-W5 to W9-11, compared to the consistent increase detected in monocultures, along with significant increases in lipofuscin. In contrast, after a significant increase in lipofuscin levels (defined as the integrated intensity from the lipofuscin pixels normalized by the number of cell pixels in a given ROI) from W3-5 to W6-8, lipofuscin levels do not increase further during W9-11 in NAM neurons. In fact, if we consider *the number* of lipofuscin pixels relative to the number of cell pixels, we observe that in the NAM neurons we start with a significantly elevated number of pixels relative to monocultures, but this number increases only insignificantly over time (Fig. S11). By comparison, an increasing proportion of lipofuscin pixels appear in the monocultures over time. This is consistent with astrocytes playing a supportive role for nearby neurons, both in terms of energy and anti-oxidant needs, consistent with previously established capability of astrocytes to supply lactate, glutamine, and glutathione (GSH) precursors to neurons [1, 51]. Monoculture neurons, on the other hand, need to maximize energy production via OXPHOS activity further contributing to ROS generation and oxidative stress (Fig. 3E-F). Collectively, these results demonstrate the ability of label-free intensity-based TPEF imaging to identify not only dynamic metabolic behavior but also interactions between different types of brain cells with significant functional implications.

### Lipofuscin autofluorescence significantly impacts optical metabolic function metrics

To demonstrate the importance of accounting for the presence of lipofuscin autofluorescence in these engineered brain tissue models, we present redox ratio and mitochondrial fragmentation assessments performed with and without the removal of lipofuscin. Given the heterogeneity in the redox ratio hues observed in Fig. 3A-C we present the redox ratio results in terms of the relative prevalence of low, mid, and high RR pixel contributions from each ROI, extracted using NNMF of the corresponding redox ratio distributions (Fig. 4A-C). Previously, we used a similar method with a three-component Gaussian distribution fit to the redox ratio distributions of neuron and astrocyte cultures treated with various concentration of Mg, and we found the prevalence of the high RR component to be an indicator of oxidative stress [54]. Consistently, neurons in monocultures have the highest High RR prevalence, with neurons in NAMs characterized by increased high RR levels compared to glial cells for all time points. Interestingly, NAM neurons show a small, insignificant drop in High RR during weeks 9-11 compared to weeks 6-8, possibly due to glial support. Another interesting observation is that monoculture neurons but not NAM neurons show an increase in Low RR components over time, indicating enhanced glycolysis or FAO along with sustained oxidative stress (Fig. 4A). In the current study, the increase in the High RR component in astrocytes (p=0.06) aligns with the increase in lipofuscin (p=0.05), suggesting higher oxidative stress in astrocytes at later weeks, also higher than in microglia (Fig. 4C). Note the dominance of the low RR pixel contributions to the glial cell redox ratio as an indicator of high levels of glycolytic and/or fatty acid oxidation activity. Overall, differences in metabolic function can be detected in a more nuanced fashion via this more detailed consideration of redox ratio, which exploits more fully the high-resolution measurements afforded by this imaging modality.

**Fig 4.**
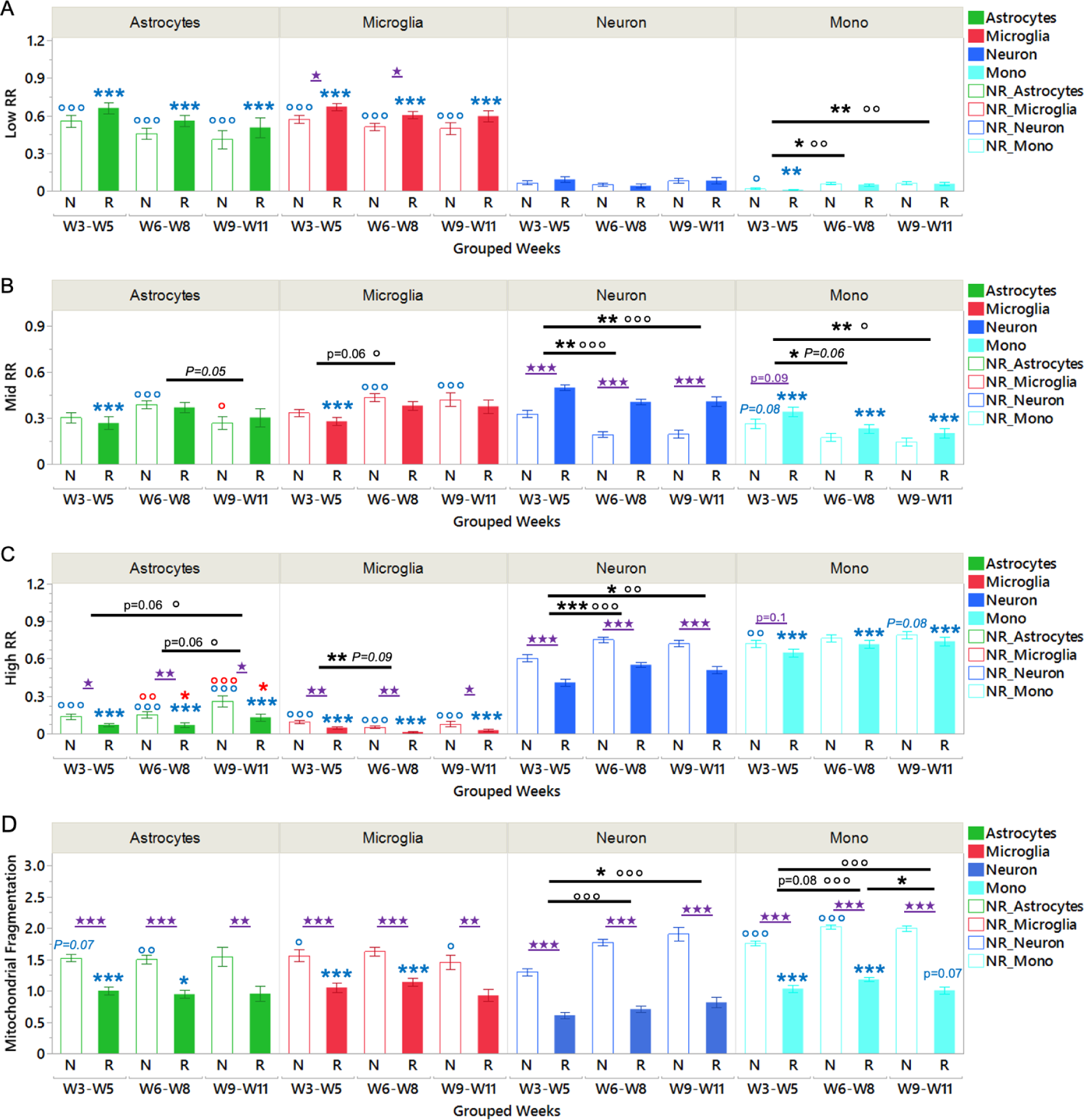
Assessment of Lipofuscin Impact on Intensity-Based Results. A. Low RR component with (R group) and without (N group) lipofuscin removal. B. Mid RR component. C. High RR component. D. Low G component. E. Mid G component. F. High G component. NR indicates groups without lipofuscin removal. Stars are the significance signs for groups with lipofuscin removal. Circles are the significance signs for groups without lipofuscin removal. Capitalized italic P values are for non-removal groups. *, **, and *** denote p < 0.05, p < 0.01, and p < 0.001, respectively. Weekly data are categorized into three ranges, with each cell type represented by a distinct color. Black asterisks indicate differences within each cell group among the three time points. Colored asterisks indicate differences among neurons, astrocytes, and microglia at a given time point (ANOVA), and between neurons in Mono and neurons in NAM.

Lipofuscin removal impacts most significantly assessments of mitochondrial fragmentation (Fig. 4D), because the bright, granule-like features of lipofuscin resemble fragmented mitochondria. Notably, neurons appear erroneously to exhibit higher levels of fragmented mitochondria compared to glial cells (Fig. 3D), which would be suggestive of enhanced glycolysis. We observe more pronounced effects following lipofuscin removal in neurons than in glial cells, because neurons contain more lipofuscin. Specifically, the Mid and High RR component values are impacted most significantly, even though differences between groups and time points remain similar for the metrics extracted with and without lipofuscin removal (Fig. 4B-C). Together, these findings highlight the significant lipofuscin contributions to human brain cell autofluorescence and the ability to exploit detailed knowledge of spectral emission from different components to mitigate its potential impact on the rigorous extraction of optical metabolic function metrics.

### Label-free fluorescence lifetime imaging is sensitive to anti-oxidant activity in human brain cells

We display representative mean lifetime-coded projections from each brain cell type in Fig. 5A-C, with bluer hues indicating decreasing mean lifetimes for all cell types over time (Fig. S12). We note, that for these assessments, we assumed a bi-exponential lifetime decay [23], which is likely an oversimplified representation of the data. However, these changes provide a qualitative representation of some of the significant phasor lifetime changes that we observe over time (Fig. S13). In an attempt to quantify these shifts, we focus on a three-component decomposition to the phasor G component distributions for each ROI, as this is the axis where we primarily detect time-dependent phasor shifts for all the cell features (Fig. S13, S14). In the hypoxia experiment we performed with microglial 2D cultures, we observe a significant decrease in the Mid G component prevalence and a significant increase in the High G component contributions (Fig. S10K-M) as OXPHOS activity becomes severely limited and is replaced by enhanced glycolysis. These results indicate sensitivity of the mid G values to the prevalence of bound NADH (which is high during high levels of OXPHOS activity) and of the low G values to the levels of free NAD(P)H (which increase as glycolysis is enhanced). The Low G component values are decreased, but not significantly upon hypoxia, but they are very different for the different types of human brain cells in our engineered brain tissue models and vary significantly over time. The data suggests that this metric may serve as a sensitive indicator of NADPH utilization by the GSH pathway to mitigate oxidative stress. This is consistent with the longer lifetime reported for bound NADPH relative to bound NADH [55].

**Fig 5.**
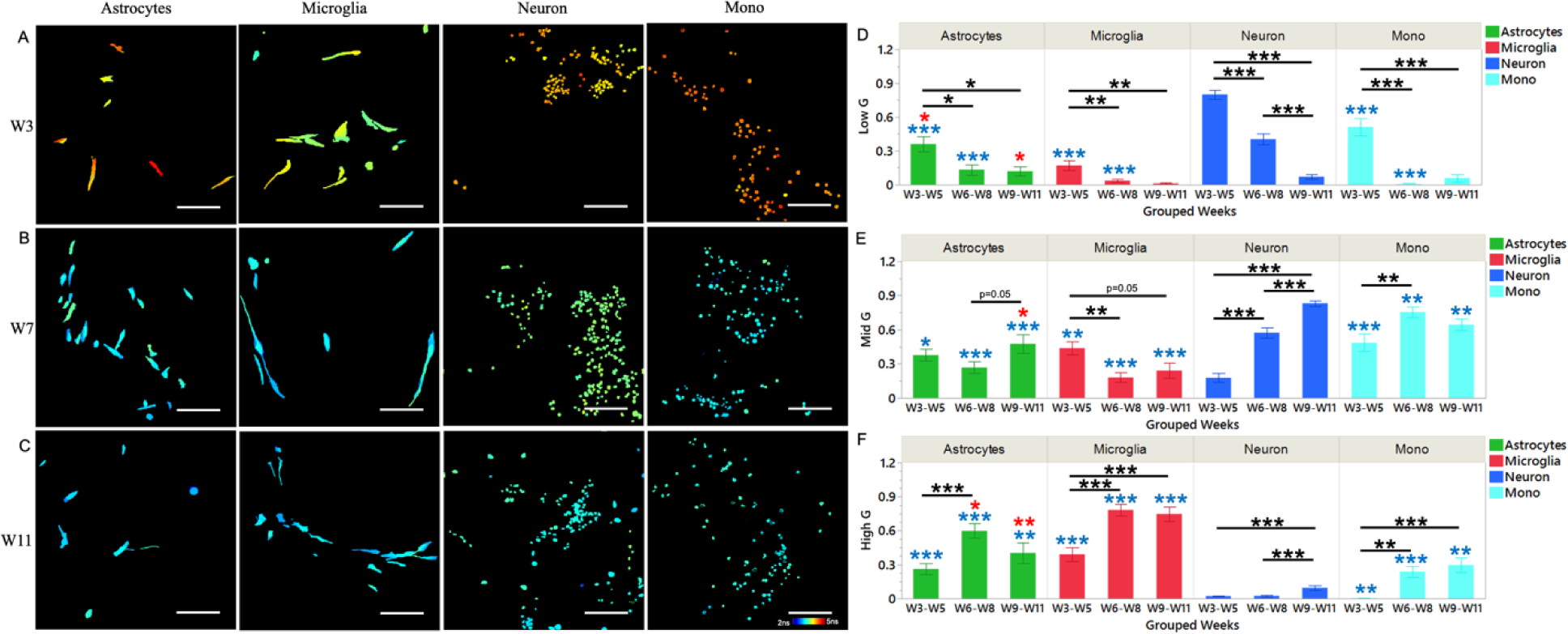
Lifetime-based NAD(P)H bounding state measurements. Representative mean lifetime color coded images for different cell populations at Week3 (A), Week7 (B), Week11 (C). (D) Low G component concentrations. (E). Mid G component concentrations. (F). High G component concentrations. (*, **, and *** denote p<0.05, p<0.01, and p<0.001, respectively). Weekly data were categorized into three ranges, each cell type represented by a distinct color. Figures in (A-C) share the same color bar and scale bar. (Black asterisks: ANOVA for each group among three time points, and colorful asterisks: Mitochondrial clustering comparisons among Neurons, astrocytes, microglia at each given group (ANOVA), and between Neurons in Mono and Neurons in NAM via pairwise two tail t-test).

All cell features show a reduction in the prevalence of phasors with Low G value coordinates. Neurons in NAMs exhibit the highest overall level of decrease in the prevalence of low G component phasors, as NADPH is being aggressively utilized to support increasing levels of OXPHOS, with limited capabilities to regenerate NADPH via the pentose phosphate pathway, possibly relying on astrocytes to replenish their NADPH pools [56]. Astrocytes exhibit higher levels of NADPH pools than microglia, especially at the initial and final time points, with astrocytes being able to maintain NADPH levels after a decrease during W6-8, consistent with their established capabilities to replenish their NADPH and maintain high anti-oxidant activity [57]. An increase in the mid G component contributions at the final time point in astrocytes is consistent with enhanced OXPHOS, also consistent with increased lipofuscin levels (Fig. 3F). Interestingly, microglia display higher levels of High G value phasors, corresponding to higher levels of free NAD(P)H, which we have associated in previous studies with higher levels of fatty acid oxidation, also consistent with lower redox ratio [23]. The abrupt drop in low G prevalence during W6-8, combined with an increase in mid and high G values, with the latter continuing to increase during W9-11 are indicative of neurons in monocultures upregulating OXPHOS more significantly than NAM neurons and then potentially engaging fatty acid oxidation to keep up with energy demands would be consistent with the lipofuscin increase and mitochondrial fragmentation decrease at the last time point (Fig 3). Overall, these results demonstrate the complementary nature of information that is provided by intensity and lifetime-based metabolic function metrics, with sensitivity to bound NADPH levels and GSH pathway activity uniquely present in lifetime measurements. They further highlight the complex and dynamic metabolic behavior of human brain cells even under standard culture conditions without any added stresses.

Finally, the lipofuscin FLIM phasor characteristics in the human brain cells of the engineered tissue models we examine are largely consistent for different cell types (Fig. S15), with neurons in monoculture having lipofuscin with a higher long lifetime than neurons in NAMs, possibly due to the different levels of GSH activity in monocultures under oxidative stress (Fig. S15). Lipofuscin phasors overlap highly with the corresponding cell phasors, and for this study we did not observe a significant impact on the derived phasor-based metrics.

### Combined use of optical metabolic function metrics enables visualization of dynamic, cell-dependent metabolic pathway utilization by different human brain cells

We use Quadratic Discriminant Analyses (QDA) to reduce the dimensionality of the 12 optical metabolic function parameters we extract from analysis of label-free, intensity and lifetime images: Mitochondrial clustering, Redox Ratio, Lipofuscin intensity, Low RR, Mid RR, High RR and NAD(P)H Short Lifetime, Long Lifetime, Bound Fraction, Low G, Mid G, High G. The results show that although all cell types exhibit time-dependent separation, astrocytes display relatively minor changes across different weeks (Fig. 6A), while other cell types show more significant differences (Fig. 6B-D). This heterogeneity suggests that astrocytes may be more able to mobilize different metabolic pathways to maintain homeostasis. When we distinguish different cell types overall, we find that astrocytes and microglia largely overlap (Fig. 6E), indicating that their metabolic functions are mostly similar in the baseline state, consistent with the indicators we observed. In contrast, neurons in monoculture and tri-culture occupied different QDA spaces, highlighting differences in pathway utilization, which were also validated by our data. These observations indicate detectable interactions between neurons and glial cells in our brain tissue model, with neurons exhibiting different dynamics under monoculture and tri-culture conditions. Overall, our baseline study suggests that this platform has significant potential in monitoring neurodegenerative diseases or injury conditions, especially by highlighting differences in metabolic pathway utilization among different cell types.

**Fig 6.**
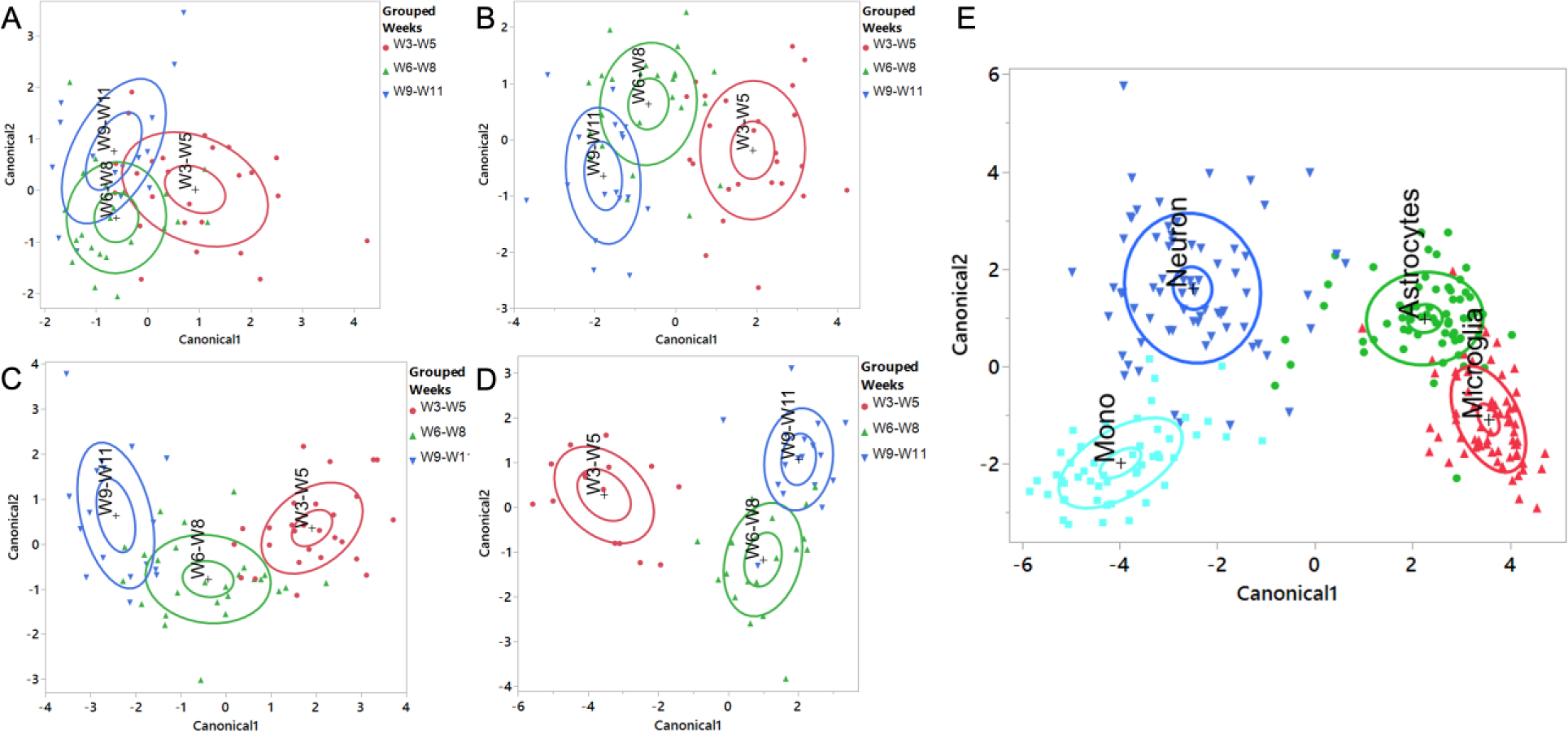
Long term dynamic monitoring of different cell types using Quadratic discriminant analyses (QDA). A combination of optical metabolic function metrics during culture, including Mitochondrial clustering, Redox Ratio, Lipofuscin intensity, Low RR, Mid RR, High RR (Derived from analysis of TPEF intensity images) and NAD(P)H: Short Lifetime, Long Lifetime, Bound Fraction, Low G AUC, Mid G AUC, High G AUC (Derived from analysis of phasor FLIM data) identify distinct dynamics and heterogeneities at multiple times over 3 months. (A-D) Canonical plots resulting from QDA using optical metabolic function metrics from Astrocytes (A) and Microglia (B), neurons at triculture (C) and monoculture (D) at different weeks. (E) Monoculture neurons are metabolically distinct from triculture Neurons, but more similar to them than to astrocytes and microglia. Astrocytes and microglia are metabolically more similar compared to neurons.

## Discussion

Label-free imaging has the potential to enable monitoring of complex cell-cell metabolic interactions that play a key role in brain function and overall organismal physiology. As a result of the need to generate energy with high efficiency, the prevalence of fluorophores that evidence the need for high anti-oxidant activity, such as NADPH, as well as the accumulation of products of the reactions with reactive oxygen species, such as lipofuscin, requires special attention in order to ensure rigorous metabolic assessments from the acquired endogenous fluorescence signals. In this study, we demonstrate the key role that spectral image acquisition and ensuing spectral decomposition play in identifying key fluorophores and corresponding spectral contributions from the endogenous TPEF emanating from human neurons, astrocytes, and microglia cultured together over a period of approximately three months in a novel silk-collagen hydrogel. Establishing the full emission spectrum of lipofuscin at two excitation wavelengths, enables us to establish a robust approach to identify lipofuscin-rich pixels from a set of images acquired only over two emission bands at these two wavelengths. Once such pixels are removed, we assess more accurately contributions form NAD(P)H and flavins and extract metrics of redox, mitochondrial fragmentation, and the NAD(P)H lifetime characteristics to define key metabolic function differences and interactions between neurons, astrocytes, and microglia.

Lipofuscin accumulation is associated with aging and oxidative stress and linked to neurodegenerative diseases like Parkinson’s and Alzheimer’s in the brain [43, 58]. This study identifies broad TPEF lipofuscin emission, excited almost equally efficiently at 755 and at 860 nm, with significant contributions almost throughout the visible spectrum, overlapping highly with emission from NAD(P)H and flavins (Fig. 2). The broad and excitation wavelength-dependent emission spectra that we observe are consistent with the presence of a combination of chemical species contributing to lipofuscin emission. This is also a key reason for variations reported in the lipofuscin fluorescence characteristics from different studies performed in different tissues [59, 60]. For example, the lipofuscin spectra features we extract are consistent with those reported from the mouse brain cortex by Eichhoff et al [61] and from isolated lipofuscin granules from retinal pigmented epithelial (RPE) cells extracted from human cadaver eyes [62]. The latter as well as several other studies highlight the dependence of the emission spectra of RPE-associated lipofuscin on the excitation wavelength and the specimen from which it is extracted [63–66]. Broad intense emission in the 430-490 nm and 570-620 nm ranges is also reported from lipofuscin deposit in the living cortex of APPswe:PS1dE9 mice, which are a popular Alzheimer’s disease model. The full emission spectrum attributed to lipofuscin from the same mouse model is shown to have a maximum at 650 nm by Chen et al [67]. A broad, but also somewhat red shifted emission with a peak at 567 nm has been reported in post-mortem frontal lobe human brains [68]. Thus, it is important to characterize the spectral excitation-emission characteristics of lipofuscin-associated fluorescence in the cell or tissue model of interest.

The lipofuscin lifetime phasor signatures are highly overlapping with those from NAD(P)H in our study (Fig. S17A), with estimated mean lifetimes of 3-3.5 ns (Fig. S17D). Reported lipofuscin lifetimes also vary significantly. In the human cortex, values in the range of 1.41-1.53 ns have been presented by Hakvoort et al [69]. Longer lipofuscin lifetimes have been demonstrated in isolated lipofuscin granules from retinal epithelial cells, as a result of oxidized products of bisretinoids [62, 70, 71]. In studies of the mouse cortex, lipofuscin TPEF associated lifetime phasors are consistent with shorter lifetimes than those extracted from NADH-associated autofluorescence contributions but appear to vary significantly even for the same mouse model [67, 72]. Given the high sensitivity of lifetime characteristics to the local environment, it is important that assessments of lipofuscin contributions be made in the samples that are being examined. While fluorescence lifetime metrics may be distinct from other cellular fluorophores in some cases, this is not necessarily always the case.

Lipofuscin isolation enables us to assess more accurately contributions from NAD(P)H and flavins, which yield important metabolic function insights. While the fluorescence intensity emission characteristics of NADH and NADPH are very similar, their physiological roles are quite distinct. NADH is key for energy production and redox regulation, including processes like glycolysis, OXPHOS and the TCA cycle [73] [74]. NADPH, on the other hand, is vital for synthesizing fatty acids, cholesterol, and steroids, as well as for antioxidant defense, nitric oxide production, and detoxification [75] [76] [77] [78] [79]. Neurons are highly vulnerable to oxidative stress due to a lower glutathione pool [80] and reduced ability to regenerate NADPH through the pentose phosphate pathway[81]. Consequently, they rely on astrocytes for glutathione transfer [82]. While we often assume NADPH fluorescence contributions to be insignificant relative to those from NADH, there are variations in NADPH content, with a mouse HPLC-MS study reporting higher NADPH/NADH concentration ratios in brain tissue compared to other tissues like the heart, liver, and skeletal muscle [83]. Further, FLIM studies reveal significant differences in their lifetimes, with bound NADPH having a longer lifetime than bound NADH [55, 75]. Our results indicate that neurons, astrocytes, and microglia exhibit a decrease in the low G component values of the lifetime phasor distributions detected at 755nm excitation and 435-585 nm emission over time in culture. This metric appears sensitive to bound NADPH fluorescence contributions (Fig. 5D) and its changes are consistent with anticipated GSH pathway activity mobilized to combat oxidative stress, as evidenced by lipofuscin accumulation and high redox ratio values. Several studies highlight the important role of astrocytes for supplying neurons with substrates such as lactate, glutamine, cysteine, and glycine [1, 51], to alleviate energy demands and to improve anti-oxidant defenses. We observe significant differences in intensity and lifetime-based metabolic function metrics, consistent with interactions between astrocytes in neurons leading to improved handling of oxidative stress relative to neurons cultured in isolation.

While bound NADH also likely influences significantly phasors with low G values (along with bound NADPH), we expect that it is a major contributor to the mid G value phasors. Significantly higher prevalence of mid G values in neurons in monocultures vs. NAM tissues support the higher levels of reliance on tricarboxylic acid-driven metabolism for energy production during the first 8 weeks in culture. However, this sort of activity is more sustainable in NAM neurons, for which mid G value phasors continue to become more prevalent during weeks 9-11. The small and insignificant increase in mid G phasor values and higher levels of high G value phasors may be indicative of enhanced fatty acid oxidation, also consistent with the increase in low RR values. The significantly higher prevalence of high G phasor values for astrocytes and microglia relative to neurons are consistent with the high glycolytic and/or fatty acid oxidation activity levels of these cells, in agreement with the RR and mitochondrial fragmentation results.

Flavins, FAD as well flavin mononucleotide (FMN), and associated flavoproteins, such as LipDH and electron-transferring flavoprotein (ETF), are also significant cellular autofluorescence contributors [33]. In studies performed with isolated mitochondria and mammalian cell models, FAD levels are significantly higher than FMN [84] [85]. In fact, LipDH is typically considered as the main source of cellular flavin signal [86] with a higher two-photon excitation action cross-section than free FAD [48]. Further, according to Kunz et al., ETF levels in mouse brain tissues are usually low, while LipDH and FAD contributions are relatively higher [87]. Our spectral decomposition identified two primary flavoprotein contributors, likely FAD and LipDH. LipDH peaks approximately at 510 nm, and its excitation efficiency at 860 nm compared to 755 nm is consistent with previous studies [48]. The FAD emission spectrum extracted from our deconvolution peaks at 560 nm, which is more red shifted than published spectra, typically is approximately at 520-540 nm [48, 87]. Although there are shape discrepancies, the epithelium-derived spectra in our study are also fit well with the brain tissue-derived shapes (Fig. S8A-D). Additional excitation wavelength and fluorescence lifetime characteristics we attribute to this component are also consistent with those of FAD.

In summary, our study reveals several important findings related to label-free, two photon metabolic function assessment of human brain cells. Firstly, the identification and removal of lipofuscin is critical for accurate metabolic analysis. Lipofuscin accumulates over time, predominantly in neurons, while the presence of glial cells can help alleviate oxidative stress in neurons. Secondly, our research highlights the value of using multimodal two-photon technologies for cell-specific metabolic analysis. These techniques allow us to accurately quantify different cellular fluorophores, specifically the activities of NADH, NADPH, and flavoproteins. Our methods can capture non-destructively and with subcellular resolution dynamic changes in the activity of key metabolic pathways, including OXPHOS, glycolysis, the glutathione pathway, and fatty acid oxidation. These studies form the foundation for further rigorous metabolic function assessments in brain tissues aiming to improve our understanding of physiological and diseased brain function or to assess the impact of novel interventions. While the engineered brain tissue models that we employ have limitations in terms of representing the complexity of human brain function, however, they also provide unique opportunities to assess the impact of well-controlled factors that are difficult to study otherwise.

## Materials and Methods

### 3D engineered brain tissue models

Neuronal monoculture (Mono) and triculture systems (NAM) are established in 3D hybrid silk-collagen hydrogel scaffolds, comprising either exclusively hiNSCs, or a combination of hiNSCs, human astrocytes, and human microglia, respectively, following previously published protocols (13, 40). mCherry-expressing HMC3 microglia were used in the TPEF intensity and FLIM imaging set of experiments, for which images were annotated. More details are provided in Supplementary Methods.

### Two photon, label-free metabolic imaging

Images are acquired using a Leica TCS SP8 microscope equipped with a tunable femtosecond laser (InSight Deep See; Spectra-Physics; Mountain View, California) that operates from 680 to 1300 nm, utilizing a water-immersion 25× objective with a numerical aperture of 0.95. Spectral images are acquired at 755 nm and 860 nm excitation within the 390-740 nm emission range, using a 10 nm bandwidth and a 10 nm step size, with a descanned HyD detector. Two-photon excitation fluorescence (TPEF) and second harmonic generation (SHG) images (1024 × 1024 pixels, 465 × 465 μm) are acquired at both 755 nm and 860 nm excitation using four non-descanned detectors, which collect light at the following emission ranges: 430 ± 12 nm for SHG imaging at 860 nm excitation, 460 ± 25 nm, 525 ± 25 nm, and 624 ± 20 nm for mCherry autofluorescence at 755 nm excitation. The incident powers for all data types are maintained at 10.9 mW for 755 nm and 19.1 mW for 860 nm throughout the experiments. Eight frame averages are acquired from each optical section at a 600 Hz scan rate. For intensity and fluorescence lifetime imaging (FLIM), the same volumes of interest are captured using four sequential 2D optical sections spaced 4 μm apart, resulting in a 465 × 465 × 12 μm³ volume. Spectral data are acquired from a single optical section within this volume. Three regions of interest (ROIs) are imaged in different areas of each scaffold to capture cellular morpho-functional heterogeneity in these tissue models. Following the acquisition of intensity data, FLIM measurements are collected at the same optical sections using 755 nm excitation, with emission collected at 460 ± 25 nm and 525 ± 25 nm with a 1-minute integration time, utilizing a Picoquant Picoharp 300 time-correlated single photon counter and SymPhoTime analysis software. Finally, spectral acquisition is performed, with two to three ROIs acquired from each scaffold, requiring 2-3 minutes to complete a full spectral acquisition per wavelength. For each type of tissue, the same three scaffolds are imaged weekly or bi-weekly for a period of 11 weeks. The microscope stage is enclosed in an incubator stage chamber maintained at 37° (Leica Microsystems) to ensure tissues are maintained at 37°C and 5% CO2 atmosphere. After each imaging session, tissues are put back to the original dish and maintained for further monitoring. Images are analyzed for data acquired from three independent monocultures and three independent NAM tissues weekly or biweekly from week 3 to week 11 in culture. In total, data are acquired from 57 ROIs from neuronal monocultures and 72 ROIs from NAM tissues.

### Spectral Analysis

Non-negative matrix factorization (NNMF) is utilized to extract cellular components from spectral shapes at both wavelengths using the “nnmf” function as detailed in the MATLAB Documentation (MathWorks, 2023). NNMF decomposes spectra into two matrices—one with the spectral features of N components and another containing the weights for each component. The microscope has a 650 nm short-pass filter to prevent the detection of 755nm excitation light. This filter introduces artifacts in the detection of emission beyond 625nm. This is not present for the spectra acquired at 860 nm excitation, because an 815 nm short pass filter is used in this case. However, to maintain consistency we spectrally deconvolve the spectra only from 390-625 nm. In total, after ISBA-based cell segmentation described in Supplemental material (SM2), we include 69 spectra from neuronal monocultures and 95 spectra from NAMs acquired from Week 2 to Week 14 of culture.

The detailed procedures we follow to identify the number of key fluorophores and their relevant contributions are outlined in Supplemental Methods.

### Two photon image analysis

Before quantifying metabolic parameters, all two-photon intensity data are subjected to a pre-processing routine to account for illumination power and detector gain calibrations. Following calibration, spectral data are processed to isolate cell locations for subsequent spectral decomposition, as detailed in Supplemental Methods. Intensity-based TPEF image analysis involves a) lipofuscin pixel identification and quantification, b) optical redox ratio assessments based on NAD(P)H and flavin contributions, and c) power-spectral density (PSD)-based analysis of the NAD(P)H intensity images to characterize the levels of mitochondrial clustering. For each 3D volume, for each cell features, after lipofuscin pixels removal, we sort all the cellular redox ratio values into a histogram ranging from 0 to 1, with a bin size as 0.1, therefore the redox distribution has an abscissa as redox ratio values, and ordinate as the frequency of each redox ratio range. The distributions of redox ratio values are further analyzed to assess relative contributions using a 3-component NNMF model to get the low, middle, and high redox components and concentrations (Fig. 1Dg). Phasor analysis involves applying cosine and sine transformations to time decay data to create g and s component maps. Plotting (g, s) pairs in phasor space allows extraction of long and short fluorescence lifetimes and the bound fraction of NAD(P)H, assuming bimodal exponential decays [23, 34]. Linear regression fits these values to a line, with intersections on the universal circle estimating free and bound lifetime components. We project these components to determine the bound fraction and mean lifetime of NAD(P)H. Notable variations in g values lead us to transform phasor g values from each ROI (four optical sections) into distributions, followed by NNMF as well as the redox ratio analysis, deriving Low G, Mid G, and High G components. Both intensity-based and FLIM-based analyses use manually annotated data to ensure accurate classification of cellular features.

### Cell feature annotations

Cell annotations are manually performed using Napari software on integrated intensity images from the 755ex/460em, 755ex/525em, 860ex/460em, and 860ex/525em detectors. In annotating neurons, we rely on their distinct round shapes. Corresponding bright field images are used to identify highly scattering features, which likely represent debris of dead or distressed microglia/astrocytes to ensure such objects are not annotated. Microglia are identified and annotated based on the presence of mCherry signals in the 755ex/624em channel, focusing on cells with distinct outlines and visible branches; Cells with ambiguous outlines are not annotated. Astrocytes are identified by their features, such as their non-round shape, the absence of mCherry signals in the 755 nm excitation/624 nm emission channel, and their size, which is larger than neurons. Neurons typically have a maximum somata size of several microns in diameter. When annotating astrocytes, care is taken to account for their proximity or overlap with microglia or other cells, especially in situations where cell outlines might be ambiguous. Analysis of the intensity and FLIM images is performed for each cell class feature identified within a given image stack separately. In monoculture, 6,261 neurons are annotated. In tricultures, 3,137 neurons, 368 astrocytes, and 587 microglia are annotated. The number of cells in each 3D volume is determined using “bwconncomp” function in MATLAB with 26-connectivity in MATLAB to identify and count the connected components representing individual cells.

### Image co-registration for 755 and 860 and for intensity and FLIM

To ensure results are calculated from the same cell volume for the two wavelengths, we determine adjustments for the focal plane differences of the objective at 755 nm and 860 nm. Specifically, for this dataset, there is a 4 μm offset in the z direction between 755 nm and 860 nm data collection, so we pair the second layer of 755 nm data with the 860 nm data. FLIM data are acquired for co-registered locations with the intensity data, but there is a system shift horizontally by 4 pixels relative to the intensity data. Therefore, we shift the annotations made based on intensity images by 4 pixels to extract the cell features for FLIM analyses.

### Lipofuscin quantification

Based on the spectral decomposition results, we examine the ratio of the pixel intensities at the 860/525 nm and 755/525 nm detectors within all annotated cells and we assign all pixels with ratios higher than 0.8 to lipofuscin. For a given ROI, we report the integrated intensity from all four images (755 nm, 860 nm excitations and 460 nm, 525 nm emissions), normalized to the corresponding number of pixels occupied by cells.

### Optical redox ratio assessments

Following lipofuscin pixel identification and removal, we consider the relative intensities of the signals detected in the 755ex/460em, 755ex/525em, 860ex/460em, and 860ex/525em detectors and we utilize the spectral shapes of NAD(P)H and flavins identified from the spectral decomposition results to quantify contributions from NAD(P)H and flavins in the imaged volumes as described in Supplemental Methods (SM7). The pixel wise redox ratio is calculated using the formula FP / (FP + NAD(P)H), where ‘FP’ includes contributions from both LipDH and FAD.

Exploiting the high spatial resolution afforded by two-photon imaging, we generate pixel-wise redox ratio maps for each cell pixel within the intensity volumes. These maps are then decomposed using NNMF into three different redox distributions that peak at ∼ 0.2, 0.45, and 0.8 redox ratio values, corresponding to low, mid, and high RRs, respectively. This segmentation utilized a methodology analogous to that demonstrated in our previous work, where the redox distribution was decomposed into 3 different gaussian components, with the higher redox component serving as a sensitive metric of oxidative stress [54]. The overall optical redox ratio is calculated from the ratio of the average FP and NAD(P)H contributions assessed from the whole field, not the pixel-wise calculations.

### PSD-based mitochondrial clustering characterization

To evaluate mitochondrial clustering, we utilize an established Fourier analysis-based approach [23, 26]. In brief, for each 3D volume, NAD(P)H images of individual cells, post-lipofuscin removal, are cloned and randomly positioned within the vacant regions of the image field to generate an image representing mitochondrial patterns across the entire field. This method, termed clone-stamping, enhances the precision of mitochondrial organization analysis by mitigating distortions introduced by sharp intensity changes at cell borders. All cell features within each 3D volume are analyzed together. We then consider the spatial frequency dependence of the squared amplitude of the 2D Fourier-transform of the image, ie the power spectral density (PSD). The segment from 0.18 μm ^−1^to the frequency corresponding to 98% of the total PSD area, as shown in Figure 1Dd, is fit using the formula *R*(*k*) = *Ak*^−*β*^ to determine the exponent’s value. Here, R represents the fit to the PSD, k is the spatial frequency magnitude, β is the power law exponent, and A is a constant. Because the clone stamping procedure is random, we perform it five times, analyze the corresponding PSD, and calculate the mean β for each image stack as a quantitative representation of mitochondrial clustering.

### Phasor fluorescence lifetime analysis

To ensure sufficient photon counts for the analysis of lifetime images, we bin the time dependent decays into 56 ranges, with each bin corresponding to 223 ps. We further implement 61x61 pixel spatial binning to aggregate photon counts from neighboring cellular pixels. Given that cells are typically spaced apart in our system, a 61x61 pixel area (55x55 μm) usually includes 1-10 distinct cells. In this manner, all spectra utilized include an average of at least 1,000 photon counts. For lipofuscin phasor calculations, all ROIs for the same scaffolds each week are integrated to ensure a sufficient number of lipofuscin pixels for line fitting (Fig. S15).

To characterize the heterogeneity within lifetime features, particularly due to variations predominantly observed along the g axis, these maps are also decomposed using NNMF into three different g components that peak at ∼ 0.34, 0.4, 0.46 g values, corresponding to low, mid, and high RRs, respectively. (Figure 1Ee).

### Statistical analysis

Pairwise two-tailed t-tests are used for comparisons between metabolic function metrics for the neurons in the monocultures and co-cultures. ANOVA followed by Tukey’s post-hoc tests are used to assess the level of significant differences among different weeks per cell type, and also among the different cell types per week. Statistical analyses are conducted using JMP 17 (SAS Institute).

### Quadratic Discriminant Analysis (QDA)

QDA is performed using the values of 12 metabolic function parameters derived from both intensity and FLIM analyses, as a means to reduce the data and visualize the metabolic states that each combination represents. The parameters include Mitochondrial clustering, Redox Ratio, Lipofuscin intensity, Low RR, Mid RR, High RR, Low G, Mid G, High G concentrations, NAD(P)H bound fraction, Long Lifetime, and Short Lifetime.

## Acknowledgments

The authors acknowledge funding from the Department of Defense Technology for the Warfighter Program (HU0001-20-2-0015), NIH R01 EB030061, NIH Research Infrastructure grant (NIH S10OD021624), NIH (P41EB027062), DoD (W911NF-23-1-0276), DoD (W81XWH2211065). Exactly that IG is a coauthor of a patent U.S. Patent No: 10,712,272, entitled System and Method for Assessing Cellular Metabolic Activity, issued on Jul 14, 2020.

## Author contributions

Conceptualization: I.G., V.L., D.L.K

Methodology: Y.Z., I.G., M.S., V.L., Y.F

Experiments: Y.Z., M.S., V.L., M.L

Cell Annotations: V.R., T.B., A.S., Y.Z., M.E.D., X.C., S.N., A.D

Data Analysis: Y.Z., I.G

Figure Preparation: Y.Z., M.S., I.G., U.O.U., Y.F.

Supervision: I.G., D.L.K., E.L.M

Writing—original draft: Y.Z., M.S., I.G

Writing—review & editing: I.G., E.L.M., D.L.K., M.S., Y.Z., V.L., U.O.U

## Data and materials availability

The key scripts used for analyses to be uploaded in the Github repository

All data needed to evaluate the conclusions in the paper are present in the paper and/or the Supplementary Materials. Raw data for this study to be deposited in online database.

## References

1. Bonvento, G. and J.P. Bolanos, Astrocyte-neuron metabolic cooperation shapes brain activity. Cell Metab, 2021. 33(8): p. 1546–1564.

2. Shan, Y., et al., Integrated Positron Emission Tomography/Magnetic Resonance Imaging for Resting-State Functional and Metabolic Imaging in Human Brain: What Is Correlated and What Is Impacted. Front Neurosci, 2022. 16: p. 824152.

3. Jamadar, S.D., et al., Task-evoked simultaneous FDG-PET and fMRI data for measurement of neural metabolism in the human visual cortex. Scientific Data, 2021. 8(1): p. 267.

4. Downes, D.P., et al., Characterization of Brain Metabolism by Nuclear Magnetic Resonance. Chemphyschem, 2019. 20(2): p. 216–230.

5. Shahsavarani, S., et al., Cortex-wide neural dynamics predict behavioral states and provide a neural basis for resting-state dynamic functional connectivity. Cell Rep, 2023. 42(6): p. 112527.

6. Guo, S., C. Zhang, and A. Le, The limitless applications of single-cell metabolomics. Current Opinion in Biotechnology, 2021. 71: p. 115–122.

7. Svatoš, A., Single-cell metabolomics comes of age: new developments in mass spectrometry profiling and imaging. Analytical Chemistry, 2011. 83(13): p. 5037–5044.

8. Benisty, H., et al., Rapid fluctuations in functional connectivity of cortical networks encode spontaneous behavior. Nat Neurosci, 2024. 27(1): p. 148–158.

9. Adler, A., et al., Sleep promotes the formation of dendritic filopodia and spines near learning-inactive existing spines. Proc Natl Acad Sci U S A, 2021. 118(50).

10. Ahn, S.J., et al., Diverse Inflammatory Response After Cerebral Microbleeds Includes Coordinated Microglial Migration and Proliferation. Stroke, 2018. 49(7): p. 1719–1726.

11. Barros, L.F., et al., Enlightening brain energy metabolism. Neurobiol Dis, 2023. 184: p. 106211.

12. Chia, T.H., et al., Multiphoton fluorescence lifetime imaging of intrinsic fluorescence in human and rat brain tissue reveals spatially distinct NADH binding. Opt Express, 2008. 16(6): p. 4237–49.

13. Kasischke, K.A., et al., Neural activity triggers neuronal oxidative metabolism followed by astrocytic glycolysis. Science, 2004. 305(5680): p. 99-103.

14. Vishwasrao, H.D., et al., Conformational dependence of intracellular NADH on metabolic state revealed by associated fluorescence anisotropy. J Biol Chem, 2005. 280(26): p. 25119–26.

15. Yaseen, M.A., et al., Fluorescence lifetime microscopy of NADH distinguishes alterations in cerebral metabolism in vivo. Biomed Opt Express, 2017. 8(5): p. 2368–2385.

16. Dora, E., et al., Effect of arterial hypoxia on the cerebrocortical redox state, vascular volume, oxygen tension, electrical activity and potassium ion concentration. Acta Physiol Acad Sci Hung, 1979. 54(4): p. 319–31.

17. Rehncrona, S., B. Chance, and G. Austin, Microheterogeneity of redox states in cerebral cortical tissue during hypoxia and ischemia. Adv Neurol, 1979. 26: p. 325–33.

18. Chance, B., et al., Intracellular oxidation-reduction states in vivo. Science, 1962. 137(3529): p. 499-508.

19. Mayevsky, A. and B. Chance, Repetitive Patterns of Metabolic Changes during Cortical Spreading Depression of Awake Rat. Brain Research, 1974. 65(3): p. 529–533.

20. Mayevsky, A. and B. Chance, Metabolic Responses of Awake Cerebral-Cortex to Anoxia Hypoxia Spreading Depression and Epileptiform Activity. Brain Research, 1975. 98(1): p. 149–165.

21. Rocheleau, J.V., W.S. Head, and D.W. Piston, Quantitative NAD(P)H/flavoprotein autofluorescence imaging reveals metabolic mechanisms of pancreatic islet pyruvate response. J Biol Chem, 2004. 279(30): p. 31780–7.

22. Shiino, A., et al., Three-dimensional redox image of the normal gerbil brain. Neuroscience, 1999. 91(4): p. 1581–5.

23. Liu, Z., et al., Mapping metabolic changes by noninvasive, multiparametric, high-resolution imaging using endogenous contrast. Sci Adv, 2018. 4(3): p. eaap9302.

24. Quinn, K.P., et al., Quantitative metabolic imaging using endogenous fluorescence to detect stem cell differentiation. Sci Rep, 2013. 3: p. 3432.

25. Varone, A., et al., Endogenous two-photon fluorescence imaging elucidates metabolic changes related to enhanced glycolysis and glutamine consumption in precancerous epithelial tissues. Cancer Res, 2014. 74(11): p. 3067–75.

26. Pouli, D., et al., Imaging mitochondrial dynamics in human skin reveals depth-dependent hypoxia and malignant potential for diagnosis. Sci Transl Med, 2016. 8(367): p. 367ra169.

27. Shiu, J., et al., Noninvasive Imaging Techniques for Monitoring Cellular Response to Treatment in Stable Vitiligo. J Invest Dermatol, 2024. 144(4): p. 912–915 e2.

28. Shiu, J., et al., Multimodal analyses of vitiligo skin identify tissue characteristics of stable disease. JCI Insight, 2022. 7(13).

29. Xylas, J., et al., Improved Fourier-based characterization of intracellular fractal features. Optics Express, 2012. 20(21): p. 23442–23455.

30. Xylas, J., et al., Noninvasive assessment of mitochondrial organization in three-dimensional tissues reveals changes associated with cancer development. Int J Cancer, 2015. 136(2): p. 322–32.

31. Pouli, D., et al., Label-free, High-Resolution Optical Metabolic Imaging of Human Cervical Precancers Reveals Potential for Intraepithelial Neoplasia Diagnosis. Cell Rep Med, 2020. 1(2).

32. Datta, R., et al., Fluorescence lifetime imaging microscopy: fundamentals and advances in instrumentation, analysis, and applications. J Biomed Opt, 2020. 25(7): p. 1–43.

33. Georgakoudi, I. and K.P. Quinn, Label-Free Optical Metabolic Imaging in Cells and Tissues. Annu Rev Biomed Eng, 2023.

34. Liu, Z., et al., Label-free, multi-parametric assessments of cell metabolism and matrix remodeling within human and early-stage murine osteoarthritic articular cartilage. Commun Biol, 2023. 6(1): p. 405.

35. Liaudanskaya, V., et al., Mitochondria dysregulation contributes to secondary neurodegeneration progression post-contusion injury in human 3D in vitro triculture brain tissue model. Cell Death Dis, 2023. 14(8): p. 496.

36. Snapper, D.M., et al., Development of a novel bioengineered 3D brain-like tissue for studying primary blast-induced traumatic brain injury. J Neurosci Res, 2023. 101(1): p. 3–19.

37. Jorfi, M., et al., Three-Dimensional Models of the Human Brain Development and Diseases. Adv Healthc Mater, 2018. 7(1).

38. Jensen, G., C. Morrill, and Y. Huang, 3D tissue engineering, an emerging technique for pharmaceutical research. Acta Pharmaceutica Sinica B, 2018. 8(5): p. 756–766.

39. Bjorklund, G.R., T.R. Anderson, and S.E. Stabenfeldt, Recent Advances in Stem Cell Therapies to Address Neuroinflammation, Stem Cell Survival, and the Need for Rehabilitative Therapies to Treat Traumatic Brain Injuries. Int J Mol Sci, 2021. 22(4).

40. Cairns, D.M., et al., A 3D human brain-like tissue model of herpes-induced Alzheimer’s disease. Sci Adv, 2020. 6(19): p. eaay8828.

41. Rouleau, N., et al., A Long-Living Bioengineered Neural Tissue Platform to Study Neurodegeneration. Macromol Biosci, 2020. 20(3): p. e2000004.

42. Lomoio, S., et al., 3D bioengineered neural tissue generated from patient-derived iPSCs mimics time-dependent phenotypes and transcriptional features of Alzheimer’s disease. Mol Psychiatry, 2023. 28(12): p. 5390–5401.

43. Moreno-Garcia, A., et al., An Overview of the Role of Lipofuscin in Age-Related Neurodegeneration. Front Neurosci, 2018. 12: p. 464.

44. Tomasi, D., G.J. Wang, and N.D. Volkow, Energetic cost of brain functional connectivity. Proc Natl Acad Sci U S A, 2013. 110(33): p. 13642–7.

45. Bélanger, M., I. Allaman, and Pierre J. Magistretti, Brain Energy Metabolism: Focus on Astrocyte-Neuron Metabolic Cooperation. Cell Metabolism, 2011. 14(6): p. 724–738.

46. Digman, M.A., et al., The phasor approach to fluorescence lifetime imaging analysis. Biophys J, 2008. 94(2): p. L14–6.

47. Zhang, Y., et al., Factors associated with obesity alter matrix remodeling in breast cancer tissues. J Biomed Opt, 2020. 25(1): p. 1–14.

48. Huang, S., A.A. Heikal, and W.W. Webb, Two-photon fluorescence spectroscopy and microscopy of NAD(P)H and flavoprotein. Biophys J, 2002. 82(5): p. 2811–25.

49. Sanchez-Hernandez, A., C.M. Polleys, and I. Georgakoudi, Formalin fixation and paraffin embedding interfere with the preservation of optical metabolic assessments based on endogenous NAD(P)H and FAD two-photon excited fluorescence. Biomed Opt Express, 2023. 14(10): p. 5238–5253.

50. Chorvat, D., Jr. and A. Chorvatova, Spectrally resolved time-correlated single photon counting: a novel approach for characterization of endogenous fluorescence in isolated cardiac myocytes. Eur Biophys J, 2006. 36(1): p. 73–83.

51. Ioannou, M.S., et al., Neuron-Astrocyte Metabolic Coupling Protects against Activity-Induced Fatty Acid Toxicity. Cell, 2019. 177(6): p. 1522–1535 e14.

52. Ghosh, S., et al., Bioenergetic regulation of microglia. Glia, 2018. 66(6): p. 1200–1212.

53. Gumbs, S.B.H., et al., Human microglial models to study HIV infection and neuropathogenesis: a literature overview and comparative analyses. J Neurovirol, 2022. 28(1): p. 64–91.

54. Stuntz, E., et al., Endogenous Two-Photon Excited Fluorescence Imaging Characterizes Neuron and Astrocyte Metabolic Responses to Manganese Toxicity. Sci Rep, 2017. 7(1): p. 1041.

55. Blacker, T.S., et al., Separating NADH and NADPH fluorescence in live cells and tissues using FLIM. Nat Commun, 2014. 5: p. 3936.

56. Jimenez-Blasco, D., et al., Astrocyte NMDA receptors’ activity sustains neuronal survival through a Cdk5-Nrf2 pathway. Cell Death Differ, 2015. 22(11): p. 1877–89.

57. Becerra-Calixto, A. and G.P. Cardona-Gomez, The Role of Astrocytes in Neuroprotection after Brain Stroke: Potential in Cell Therapy. Front Mol Neurosci, 2017. 10: p. 88.

58. Brunk, U.T. and A. Terman, Lipofuscin: mechanisms of age-related accumulation and influence on cell function. Free Radic Biol Med, 2002. 33(5): p. 611–9.

59. Różanowska, M.B., *Lipofuscin, Its Origin, Properties,* and Contribution to Retinal Fluorescence as a Potential Biomarker of Oxidative Damage to the Retina. Antioxidants (Basel), 2023. 12(12).

60. Wolf, G., Lipofuscin, the age pigment. Nutr Rev, 1993. 51(7): p. 205–6.

61. Eichhoff, G., et al., In Vivo Ca2+ Imaging of the Living Brain Using Multi-cell Bolus Loading Technique, in Calcium Measurement Methods, A. Verkhratsky and O.H. Petersen, Editors. 2010, Humana Press: Totowa, NJ. p. 205-220.

62. Różanowska, M.B. and B. Różanowski, Photodegradation of Lipofuscin in Suspension and in ARPE-19 Cells and the Similarity of Fluorescence of the Photodegradation Product with Oxidized Docosahexaenoate. Int J Mol Sci, 2022. 23(2).

63. Palczewska, G., et al., Noninvasive multiphoton fluorescence microscopy resolves retinol and retinal condensation products in mouse eyes. Nat Med, 2010. 16(12): p. 1444–9.

64. Palczewska, G., M. Wojtkowski, and K. Palczewski, From mouse to human: Accessing the biochemistry of vision in vivo by two-photon excitation. Prog Retin Eye Res, 2023. 93: p. 101170.

65. Feldman, T.B., et al., Fluorescence emission and excitation spectra of fluorophores of lipofuscin granules isolated from retinal pigment epithelium of human cadaver eyes. Russian Chemical Bulletin, 2010. 59(1): p. 276–283.

66. Haralampus-Grynaviski, N.M., et al., Spectroscopic and morphological studies of human retinal lipofuscin granules. Proceedings of the National Academy of Sciences, 2003. 100(6): p. 3179–3184.

67. Chen, C., et al., In Vivo Near-Infrared Two-Photon Imaging of Amyloid Plaques in Deep Brain of Alzheimer’s Disease Mouse Model. ACS Chemical Neuroscience, 2018. 9(12): p. 3128–3136.

68. Lochocki, B., et al., Multimodal, label-free fluorescence and Raman imaging of amyloid deposits in snap-frozen Alzheimer’s disease human brain tissue. Communications Biology, 2021. 4(1): p. 474.

69. Hakvoort, K., et al., Shedding light on human cerebral lipofuscin: An explorative study on identification and quantification. J Comp Neurol, 2021. 529(3): p. 605–615.

70. Semenov, A.N., et al., Protein-Mediated Carotenoid Delivery Suppresses the Photoinducible Oxidation of Lipofuscin in Retinal Pigment Epithelial Cells. Antioxidants, 2023. 12(2): p. 413.

71. Yakovleva, M.A., et al., Fluorescence characteristics of lipofuscin fluorophores from human retinal pigment epithelium. Photochemical & Photobiological Sciences, 2020. 19(7): p. 920–930.

72. Gomez, C.A., et al., Cerebral metabolism in a mouse model of Alzheimer’s disease characterized by two-photon fluorescence lifetime microscopy of intrinsic NADH. Neurophotonics, 2018. 5(4): p. 045008.

73. Xiao, W., et al., NAD(H) and NADP(H) Redox Couples and Cellular Energy Metabolism. Antioxid Redox Signal, 2018. 28(3): p. 251–272.

74. Imai, S.I. and L. Guarente, It takes two to tango: NAD(+) and sirtuins in aging/longevity control. NPJ Aging Mech Dis, 2016. 2: p. 16017.

75. Bonora, M., et al., A mitochondrial NADPH-cholesterol axis regulates extracellular vesicle biogenesis to support hematopoietic stem cell fate. Cell Stem Cell, 2024. 31(3): p. 359–377.e10.

76. Frederiks, W.M., et al., NADPH production by the pentose phosphate pathway in the zona fasciculata of rat adrenal gland. J Histochem Cytochem, 2007. 55(9): p. 975–80.

77. Förstermann, U. and W.C. Sessa, Nitric oxide synthases: regulation and function. Eur Heart J, 2012. 33(7): p. 829–37, 837a-837d.

78. Hamdane, D., et al., Structure and function of an NADPH-cytochrome P450 oxidoreductase in an open conformation capable of reducing cytochrome P450. J Biol Chem, 2009. 284(17): p. 11374–84.

79. Wang, X.-x., et al., NADPH is superior to NADH or edaravone in ameliorating metabolic disturbance and brain injury in ischemic stroke. Acta Pharmacologica Sinica, 2022. 43(3): p. 529–540.

80. Raps, S.P., et al., Glutathione is present in high concentrations in cultured astrocytes but not in cultured neurons. Brain Res, 1989. 493(2): p. 398–401.

81. Bolaños, J.P. and A. Almeida, The pentose-phosphate pathway in neuronal survival against nitrosative stress. IUBMB Life, 2010. 62(1): p. 14–8.

82. Pérez-Sala, D. and M.A. Pajares, Appraising the Role of Astrocytes as Suppliers of Neuronal Glutathione Precursors. International Journal of Molecular Sciences, 2023. 24(9): p. 8059.

83. Lu, W., et al., Extraction and Quantitation of Nicotinamide Adenine Dinucleotide Redox Cofactors. Antioxid Redox Signal, 2018. 28(3): p. 167–179.

84. Gianazza, E., et al., Coordinated and reversible reduction of enzymes involved in terminal oxidative metabolism in skeletal muscle mitochondria from a riboflavin-responsive, multiple acyl-CoA dehydrogenase deficiency patient. Electrophoresis, 2006. 27(5-6): p. 1182–98.

85. Hühner, J., et al., Quantification of riboflavin, flavin mononucleotide, and flavin adenine dinucleotide in mammalian model cells by CE with LED-induced fluorescence detection. Electrophoresis, 2015. 36(4): p. 518–25.

86. Zou, Y., et al., Analysis of redox landscapes and dynamics in living cells and in vivo using genetically encoded fluorescent sensors. Nat Protoc, 2018. 13(10): p. 2362–2386.

87. Kunz, W.S. and F.N. Gellerich, Quantification of the content of fluorescent flavoproteins in mitochondria from liver, kidney cortex, skeletal muscle, and brain. Biochem Med Metab Biol, 1993. 50(1): p. 103–10.

