## Supplementals for "Multi-modal, Label-free, Optical Mapping of Cellular Metabolic Function and Oxidative Stress in 3D Engineered Brain Tissue Models"

Yang Zhang *et al.*

**The PDF includes:**

Supplementary Methods SM1-SM8

- SM1. 3D In Vitro Human Monoculture and Triculture Brain Tissue Model Fabrication: Viability staining
- SM2. Image Segmentation Baseline Algorithm (ISBA) for cell feature extraction:
- SM3. Spectral decomposition
- SM4. Epithelium Tissue Collection and Imaging
- SM5. Lipofuscin accumulation induction and imaging in HMC3 cell cultures
- SM6. Lipofuscin removal for FLIM data
- SM7. Crosstalk Adjustment
- SM8. Hypoxia experiment to validate optical metabolic function metric sensitivity

Supplementary Appendix 1

Supplementary Figures S1-S17

Supplementary References

**Supplementary Methods**

***SM1. 3D In Vitro Human Monoculture and Triculture Brain Tissue Model Fabrication:***

Neuronal monoculture (Mono) and triculture systems (NAM) are established in 3D silk-collagen hydrogels, comprising either human induced neural stem cells (hiNSCs) or a combination of hiNSCs, human astrocytes, and microglia, respectively, following a previously published protocol [1]. Human induced neural stem cells (hiNSCs) (127), human primary astrocytes (Sciencell Research Laboratories, cat. no. 1800), and HMC3 human microglial cells (ATCC, cat. no. CRL-3304) are allowed to reach nearly 100% confluency on the day of triculture seeding. Silk scaffolds freshly coated with poly-ornithine (PLO) (Sigma, cat.no. A004C) and laminin are placed in 96-well plates, and a vacuum manifold is used to remove excess liquid (128). Neuronal monocultures are embedded with 1 million hiNSCs per tissue, while for the tricultures, each scaffold is seeded with a 40μL cell suspension containing 2:0.5:0.1 million hiNSCs, astrocytes, and microglia, respectively, followed by 30 minutes of incubation at 37°C to allow cell adhesion to the scaffolds. To ensure efficient cell attachment, the freshly seeded scaffolds are submerged in 150μL of media composed of NeuroBasal medium supplemented with 2% B-27, 1% Anti-Anti, 1% Glutamax, all from Invitrogen. The media is supplemented with 1% astrocyte growth factors (Sciencell Research Laboratories) for the tricultures. The cell-seeded scaffolds are incubated overnight in a tissue culture incubator (37°C, 5% CO2 in a humidified atmosphere), and the following day, they are transferred to new 96-well plates to remove the unattached cells, embedded with 100μL of collagen type I solution (Corning or R&D systems, 3 mg/mL with a pH adjusted to 7.0-7.2 with NaOH), and incubated for 30 minutes at 37°C to allow the collagen gel to crosslink. Media (150μL) is added to all scaffolds and incubated for 24h at 37°C. The following day, 3D tricultures are moved into 48-well plates with 1 ml of media in each well. Half the media volume is changed every fourth day until the end of the experiments.

***SM2. Image Segmentation Baseline Algorithm (ISBA) for cell feature extraction:***

The Image Segmentation Baseline Algorithm (ISBA) is employed for extracting cell features from autofluorescence and SHG signals. Integrated emission images at 755nm and 860nm excitation display autofluorescence from silk, collagen gel, and cells, while the SHG signal from collagen fibers is captured at 860nm excitation/430nm emission (Fig. S1A-C). For each emission stack, the third dimension is lambda, comprising 34 images ranging from 390 nm to 740 nm emission, each with a 10 nm range and step size, collected at the same excitation wavelengths and ROI. Strong SHG signals are associated with strong collagen crosslink associated TPEF signals [2], highlighting the challenge of using intensity thresholding alone for cellular feature extraction. Due to its broad excitation and emission range, collagen autofluorescence significantly contributes to the signal in label-free images, making it essential to distinguish collagen-rich regions (SHG+) from collagen-free areas (SHG-) for accurate cell identification (Fig. S1D). Each region of interest (ROI) is manually outlined to exclude silk fragments, ensuring a focus on cell-rich areas of interest (Fig. S1E). For images acquired with the non-descanned detectors, we use the integrated signals from the two detectors centered at 460 and 525 nm at each excitation wavelength.

The ISBA, as detailed in Supplementary Appendix 1, processes 2D/3D images to generate cell masks by exploiting information captured at different excitation-emission wavelength images, including SHG signals. For each excitation wavelength (755nm or 860nm), ISBA computes the sum of 460nm and 525nm emission wavelengths (or the integrated spectral intensity image) and applies a 3x3 average filter to reduce intensity variations between neighboring pixels. Silk and near-silk regions are delineated from the filtered image by multiplying with a manually generated silk-exclusion mask. To isolate SHG+ pixels, a three-level Otsu threshold is applied to the SHG channel. Values below the first threshold level identify as SHG-, while pixels above the second threshold level correspond to SHG+ locations with strong crosslinking signals. The SHG+ pixels are used to assess the colocalized crosslinking intensity level in the summed 460nm and 525nm images for each wavelength. Before quantifying the crosslinking intensity levels, three experimentally determined thresholds (as described in steps 8-10) are applied to identify and exclude non-cellular bright spots, which occasionally appear in ROIs and have intensities several times higher than cell features. Subsequently, for each excitation wavelength, the crosslinking intensity levels are determined separately for SHG+ and SHG- locations. This process excludes all pixels below the determined crosslinking intensity threshold to retain only cell features, under the assumption that cell features exhibit higher intensity than the background surrounding them. The final cell mask generated by Algorithm 1 has the same dimension as the input data, derived by combining the cell masks determined at the SHG+ and SHG- locations, as well as those of the bright spots identified at each excitation level. Notably, each slice of the final cell mask depicts cellular locations at different depths (if more than 1 optical sections along z depth).

To assess the accuracy of the ISBA algorithm to identify cell objects, we compare its performance to manual annotations of neurons and astrocytes (non-descanned images, all in 3D). As shown by the overlay images of 755 nm/460 nm and 860 nm/525 nm channels, glial cells (astrocytes and microglia) relatively have stronger NAD(P)H signals and weaker flavin signals, appearing green, while neurons appear magenta or white due to similar signal intensities in both channels (Fig. S2A-B). Therefore, the cell features identified in 755 nm data contain both neurons and glial cells, but the cell features identified in 860 nm data only show neurons as the glial flavin signals are lower than or similar to the crosslinking signals so that get removed. Thus, we assign the cell pixels identified at 860 nm as neurons, while the pixels absent from 860 nm cell masks but present in 755 nm masks are considered glial pixels. As illustrated in Fig. S2C-D, the maximum projections present neurons as green and glial cells as magenta. White pixels indicate overlapping signals at different z-depths, but at a single depth, there are no overlapping pixels. To assess the performance of ISBA we compare the number of objects it identifies to the manual annotations, both in terms of the number of objects as well as the functional metabolic readouts (redox ratio) associated with the objects. ISBA identifies approximately 93% of annotated neurons in monocultures and 90% annotated neurons in tricultures (Fig. S3D-E), reported at the ISBA efficiency. However, it also identifies significantly more objects than the available annotations, as indicated by the overall identification efficiency, defined as the number of objects identified by ISBA or annotations over the integrated number of all objects identified by either approach. Importantly, the redox ratio readouts of all the annotated and ISBA segmented neuronal features are nearly identical, indicating that the segmented features are likely neurons or neuronal axon segments that have not been annotated. However, the ISBA efficiency is only 46% for annotated astrocytes (Fig. S3F), even though the overall number of objects it identifies is still higher than that from annotations. The redox ratio of the ISBA identified astrocytes tends to be lower than that of the annotated astrocytes. This indicates that ISBA is less sensitive to astrocytes that may have lower NAD(P)H intensities and thus higher redox ratio values. This is reasonable, since we established ISBA as a conservative approach to limit potential detection of collagen associated fluorescence. As a result, sensitivity to weakly autofluorescent cells, especially within a collagen rich portion of the scaffold may be missed. Nevertheless, we note that this limitation does not impact our ability to detect metabolic differences between the W3-5 and W6-8 time points. Especially for this study, where the ISBA algorithm is used only for cell segmentation in the spectral image analysis, these results demonstrate that the cell features identified represent the metabolic heterogeneity of the annotated cells.

***SM3. Spectral decomposition***

To identify the spectral signatures of key contributors to the fluorescence spectra acquired from neuronal monocultures and NAM engineered brain tissue models over three months in culture we relied on non-negative matrix factorization, using the nnmf.m function in Matlab (2023b). The set of acquired spectra and the number of components that are expected to contribute to the spectra are provided as inputs. The function provides the spectral shapes and relative weights of each component that yield a minimum root mean square residual. The shapes of the component spectra will not have negative values, unlike the components derived from principal component analysis. If information is known about the shapes and/or weights of any of the components, they can be entered as constraints helping to ensure that the extracted spectral components are physiologically meaningful. It is well established that NAD(P)H and flavins contribute to the two-photon excited autofluorescence of all cells at the excitation wavelengths we considered in this study. Since NAD(P)H is the best defined chemically component, we performed measurements in immature epithelial cells, known to have high levels of NADH TPEF validated by mass spectrometry measurements [3]. These measurements (see SM4 below) led to the definition of the NAD(P)H spectrum, which was used as one of the fixed components for the spectral decomposition of the engineered brain tissue spectra. We assume that NAD(P)H spectral contributions are not significant at 860 nm excitation [4]. A three-component fit, with the NAD(P)H spectrum as one of these components, does not appear to capture all variations in the data, as shown by distinct peaks in the residual map (Fig. S5a). The residual map improves with a four-component fit (Fig. S5B,b), but no further improvement or additional component shapes are observed with a five-component fit (Fig. S5C, c). We thus focus on ensuring that the four components extracted from this process have biophysically meaningful features. For fluorophores, such as NAD(P)H or specific types of flavins, like FAD and LipDH, we expect that the emission spectrum will change only in intensity but not in shape as a function of excitation wavelength. This means the brightness may vary, but the spectral profile, including peak position, remains consistent. We perform decomposition on the concatenated emission spectra by appending the 860 nm data points after the 755 nm data points, forming a single continuous spectrum. This method ensures that the analysis considers both excitation wavelengths in a unified manner. However, the nnmf function in MATLAB does not provide a constraint to use identical spectral shapes for both the 755 nm and 860 nm data, which means the resulting components may differ across these wavelengths. We thus fix the shapes of two of the four components that are derived to describe the acquired spectra at 860 nm excitation (in reality three, since NAD(P)H does not contribute at 860 nm excitation) (Fig. S5B). We smooth these two shapes extracted from the initial four component decomposition for the 860 nm excitation wavelengths and reproduced the shapes for 755 nm excitation, using the relative excitation efficiency of the original decomposition, which is consistent with those reported for LipDH and FAD [4], even though the emission of the component with a 560 nm peak is more red-shifted. A final decomposition is performed with fixed shapes for NAD(P)H, LipDH and FAD to extract contributions from a fourth component, which is efficiently excited at both wavelengths, and has a broad emission which is a little different for 755 nm and 860 nm excitation (Fig. S5D). This is not unreasonable for fluorophores, such as lipofuscin, that are actually complex mixtures of fluorescing components.

To validate the component identities, we perform a number of supportive experiments. For example, we find that the brain cell-derived shapes (Fig. S5D) fit well spectra acquired from epithelial tissues (Fig. S6A-D). Mature epithelia show some lipofuscin presence, unlike immature epithelia, which also contain both flavin proteins. Previous studies report that LipDH has lifetimes of 0.28 ns, 0.88 ns, and 4.14 ns, while FAD has a free form lifetime of 2.47 ns, which quenches rapidly when bound. Given the emission profiles (Fig. 2C-D), LipDH contributes more to the 460EM channel, while FAD’s contribution increases significantly at 525em. The phasor lifetime differences detected at the two emission bands and at 860 and 910 nm excitation are consistent with the attribution of the derived components with peaks at 510 and 560 nm to LipDH and FAD (Fig. S6E-I).

***SM4. Epithelium Tissue Collection and Imaging:***

Rat cheek tissue samples are excised immediately following euthanasia and placed in clean 35mm glass-bottom dishes with a drop of PBS to prevent drying during imaging. Spectral data are collected from three locations in immature (Fig. S4A) and mature (Fig. S4C) epithelial areas at a depth of approximately 30-40 μm below the surface. For each ROI, full spectral images are acquired at 755 nm and 860 nm excitation wavelengths within the 390-740 nm emission range, with a 20 nm bandwidth and 10 nm step size (512×512 pixels, 290.6×290.6 μm, 40X water immersion objective, N.A. = 1.1). Simultaneously, fluorescence intensity and lifetime data are collected at the same locations using 755 nm, 860 nm, and 910 nm wavelengths, with a FLIM integration time of 1 minute.

Epithelial tissue images are segmented using a three-level Otsu threshold to identify cellular pixels, with those above the first threshold considered as such (Fig. S4B, D). The average cell spectrum intensity at both 755 nm and 860 nm excitation is calculated by dividing the total fluorescence yield by the number of cell pixels (Fig. S4E, F). At 755 nm excitation, the emission spectra show a sharp drop-off at around 660 nm due to the short pass filter, which is designed to block wavelengths longer than a specified cutoff, excluding the excitation light and preventing it from interfering with the emission measurements. After capturing the spectral shapes, an interpolant function extrapolates the spectral data to a 5 nm step size, concatenating the two excitation wavelength spectra from 390-625 nm. Both two-component and three-component non-negative matrix factorization (NNMF) result in a consistent NAD(P)H shape, with the three-component analysis providing a better fit (0.007 vs. 0.005). Significant overlap is observed in the two flavin shapes (Fig. S4G).

In our previous studies, NAD(P)H and flavin proteins are identified as the main fluorophores in epithelial tissues [5]. Undifferentiated epithelial cells, smaller and more proliferative than mature cells, exhibit high levels of glycolysis, resulting in elevated NAD(P)H levels in the cytoplasm. This is evident in Fig. S4A, E, where the TPEF signal at 755 nm excitation is significantly higher than at 860 nm (Green). The low 860 nm signal indicates low flavin protein concentration, consistent with the low redox state of proliferative cells. Conversely, mature epithelial cells exhibit higher oxidative phosphorylation (OXPHOS) levels and a higher redox ratio, resulting in increased flavin protein TPEF signals (Fig. S4C, F). The consistent NAD(P)H shape is used as a component for the spectral deconvolution of engineered brain tissue spectra, extracting three additional components, two of which align with LipDH and FAD emission characteristics. The deconvolution, performed on concatenated 755 nm and 860 nm spectra, reveals significantly higher intensities at 755 nm (Fig. S5A-D).

***SM5. Lipofuscin accumulation induction and imaging in HMC3 cell cultures***

The HMC3 microglia cell line (sourced from ATCC) is cultured in DMEM medium (ATCC) supplemented with 10% fetal bovine serum (Invitrogen) and 1% Anti-Anti (Invitrogen) in a 37°C, 5% CO2 incubator. The medium is changed every three days. When the cells reach 70-80% confluence, they are trypsinized with 0.25% Trypsin/EDTA at 37°C for 3 minutes. The trypsin activity is quenched with complete medium, and the cells are centrifuged at 1000 rpm for 5 minutes. The cell pellet is resuspended in complete medium for further experiments. To study lipofuscin under oxidative stress conditions, we treat the cells with lipopolysaccharide (LPS). LPS is known to stimulate inflammatory response in microglia, leading to increased levels of reactive oxygen species (ROS). In this experiment, 100,000 HMC3 cells are placed in wells of a 12-well glass-bottom plate. The cells are treated with 100 ng/ml of LPS for 24 hours to induce inflammation and quantify lipofuscin production. Controls are treated with PBS, the same diluent for LPS.

Control and LPS-treated HMC3 cells are imaged using a 40x water immersion objective (N.A. 1.1) to achieve high-resolution images of cellular features. Two-dimensional intensity and spectral autofluorescence images are acquired 24 hours post-LPS treatment at both 755 nm and 860 nm wavelengths, using a 512 x 512 pixels format with a physical size of 290.6 x 290.6 μm. The illumination powers are set to 14.1 mW and 24.7 mW for 755 and 860 nm excitation, respectively. At least one spectrum is acquired at each wavelength for each well using a descanned HyD detector. 2 control and LPS-treated spectra are acquired from 2 independent samples. In total, 14 different ROIs are acquired from the control and different LPS treated groups. In the experiment (using the finalized LPS concentration/incubation time), following spectral acquisition, a tile scan capturing at least 20 fields of intensity images at 390-740 nm emission is performed. Imaging locations are marked with four markers at the bottom of the glass dish before imaging.

Following two photon imaging, untreated and LPS-treated HMC3 cells were incubated with Nile blue A for lipofuscin granules identification via confocal imaging. Specifically, the cell medium is removed from the cells, and the cells are washed three times with 2 ml PBS. The PBS is then discarded, and the cells are fixed with 4% paraformaldehyde in PBS for 15 minutes at 37°C. After discarding the fixation buffer, the cells are washed twice with 2 ml PBS for 2 minutes each and incubated in 2 ml pre-warmed medium with 200 nM Nile Blue A for 15 minutes at 37°C. Subsequently, the incubation medium is discarded, and the cells are washed with 2 ml PBS for 2 minutes. Finally, 2 ml pre-warmed medium is added, and the cells are examined under a confocal microscope for lipofuscin bright spots. Confocal images are acquired post-fixation and staining using a HyD detector in photon counting mode, with an 8-frame accumulation. Briefly, the process followed was: Before staining with Nile Blue A, the entire area was imaged using tile scans, acquiring more than 20 ROIs for autofluorescence TPEF. This allowed for the colocalization of areas of interest before and after staining and identification of the exact same cell features. After fixation and staining, confocal images were acquired from the same locations. Additionally, a 3x zoomed image from the same field was obtained with some z depth adjustment to see the lipofuscin granules better. Each sample was imaged at a minimum of three different locations. The Nile Blue A signal is excited at 638 nm with emission collected at 650-700 nm. Two rounds of LPS studies are conducted, each containing two wells, yielding consistent results, and all two-photon images are calibrated for illumination power and gain.

To achieve the lipofuscin spectra (Fig. S7), lipofuscin pixels are isolated by using integrated emission images per wavelength from 14 different ROIs. For each ROI, the spectral images starting from 500 nm emission at 755 nm excitation (to avoid too much NAD(P)H fluorescence), and the full range of 860 nm images (34 emission images) are integrated separately as two 2D image. To ensure the same threshold is applied to all conditions, all generated 2D images are stitched together to form a large image for each wavelength. Threshold is customized based on a three-level Otsu’s threshold for the cytoplasm and lipofuscin granules. (Pixels above 6% of the first Otsu’s threshold are considered as cells, including lipofuscin granules, while pixels above 50% of the first threshold represent bright lipofuscin granules). This is to make sure the observed lipofuscin features can be identified with the conservative, and same threshold for every spectral ROI to extract the mean spectra at different excitations.

To isolate lipofuscin granules from non-descanned two-photon images acquired during LPS treatment optimization (different concentrations/incubation times leads to different intensity level in lipofuscin), the integrated image from all the ROIs with all the conditions (per excitation wavelength, sum the 460em and 525em image) are tiled into a single ROI. For lipofuscin granule identification, a three-level Otsu’s threshold is applied to the tiled image to separate pixels into three classes: background, cytoplasm, and lipofuscin granules. During LPS treatment optimization, we rely on the signal to noise ratio (SNR), calculated by comparing the intensity ratio of mean lipofuscin intensity with the mean cytoplasmic intensity. Cells are identified using a global Otsu’s threshold on each integrated image separately, with pixels above the threshold considered as cells. To fully remove lipofuscin pixels from cytoplasmic intensity calculations, pixels above 25% of the Otsu’s threshold are removed as lipofuscin, then mean lipofuscin pixel ratio are calculated (Fig. S7I).

To optimize the Nile blue A staining protocol, after staining and confocal imaging, to isolate and quantify the SNR, again by comparing the intensity ratio of mean lipofuscin staining intensity with the mean cytoplasmic intensity, each confocal image is normalized to a range of 0 to 1. Cell identification employs a global Otsu’s threshold, classifying pixels above the threshold as cells. To further distinguish lipofuscin granules from cellular pixels, a three-level Otsu’s threshold is applied, segmenting pixels into background (medium), cytoplasm, and lipofuscin granules, the pixels with value above the 2^nd^ level are lipofuscin.

***SM6. Lipofuscin removal for FLIM data***

For FLIM data, collected only at 755 nm, images are power and gain calibrated. Co-registration of FLIM intensity images ensures accurate lipofuscin identification by examining the 755ex/460em to 755ex/525em intensity ratios. Ratios between 0.5 and 0.8 are indicative of lipofuscin, aligning with specific emission area-under-curve (AUC) ratios for lipofuscin, NADH, LipDH, and FAD (Fig. 2C-D). To achieve accurate identification, a horizontal shift of 4 pixels is applied to co-register the cell and lipofuscin masks due to an offset between decay matrices and FLIM intensity data. This method successfully isolates lipofuscin, allowing us to obtain its emission spectra from both 2D microglia and 3D cultures, which closely match lipofuscin shapes obtained from spectral deconvolution (Fig. S7K-L).

***SM7. Crosstalk Adjustment:***

To accurately estimate NAD(P)H levels from optical redox calculations, it is useful to quantify and eliminate the potential contributions of LipDH and FAD in the NAD(P)H channel (755ex/460em, denoted as I74). Expanding on previous methods [6], we propose a model assuming the total spectral emission from a sample equals the linear sum of contributions from different fluorophores at that wavelength setting (Equations 1-3). In this model, capital letters A, B, and C represent the NAD(P)H fluorescence in the I74 channel, and the LipDH and FAD fluorescence intensities in the I85 channel (860ex/525em), respectively. Equation 8 expresses I75 (755ex/525em) as a combination of NAD(P)H, LipDH, and FAD, adjusted for detection efficiency differences. The parameters ξ4 and ξ5 represent the detection efficiencies of the two detection arms, with ξ4/ξ5 indicating the relative performance of the detectors, consistently measured at 0.71 when comparing spectra and relative intensities collected by the two detectors for NADH (Equations 4) and riboflavin solutions (Equations 5).

In this framework, $\alpha_{4}$and $\alpha_{5}$ are the area under the curve (AUC) of NAD(P)H in the 460em and 525em detection ranges, respectively (Fig. S8). Similarly, $\beta_{4}$ and $\beta_{5}$ represent the AUC of LipDH, while $\gamma_{4}$ and $\gamma_{5}$ represent the AUC of FAD in each detection range. $Eff_{LipDH}$ and $Eff_{FAD}$ indicate the excitation efficiencies of LipDH and FAD at 755ex and 860ex (based on the spectral peak ratios in Fig. 2C). By solving Equations 1-3, we can calculate the parameter A, which represents NAD(P)H amount in the 755ex/460em channel without flavin crosstalk, enabling more accurate redox ratio calculations. This crosstalk removal method establishes a foundation for precise redox ratio assessments in current and future experiments.

To verify the efficacy of our crosstalk removal technique from non-descanned images, we compare redox ratio values derived from deconvolution concentrations and crosstalk-adjusted intensity images (Fig. S9). Both spectral and intensity data yielded similar redox ratio results, with 2D spectral data processed using ISBA and intensity redox ratios based on 3D volume annotation.

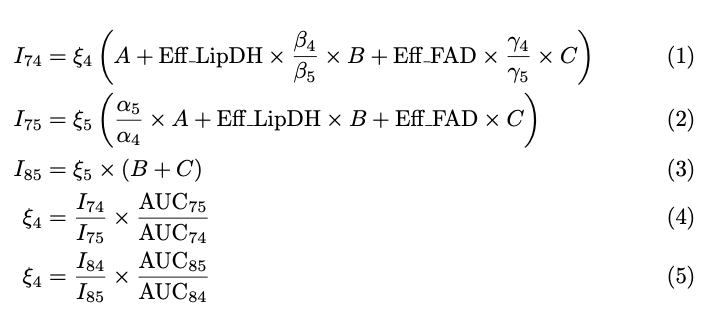

***SM8. Hypoxia experiment to validate optical metabolic function metric sensitivity:***

To validate the sensitivity of key metabolic function metrics extracted from analysis of label-free TPEF images, we perform a set of hypoxia induction experiments, since hypoxia is known to inhibit OXPHOS activity, leading to increases in NAD(P)H TPEF (since it is not utilized as efficiently for ATP production), and mitochondrial fragmentation, along with decreases in the fraction of NAD(P)H present in bound form in the mitochondria [7, 8]. To ensure the stability of the imaging dish during the experiment, we 3D print a plate holder (Fig. S10A). The experiment requires inserting a temperature sensor, an oxygen sensor, and a nitrogen needle into the medium of the imaging dish through an open hole at the cap. The nitrogen needle remains in place throughout the hypoxia and imaging periods to expel oxygen while retaining the culture medium. The cap hole is created by removing the glass bottom from a 35 mm glass-bottom imaging dish (Fig. S10B). The cap is then sealed with the dish containing cell cultures, ensuring a complete seal, including the cap area (Fig. S10C), during hypoxia. The three components protrude through the hole, as highlighted by the arrows in Fig. S10B-C.100,000 HMC3 cells are seeded in each of three 35 mm glass-bottom culture dishes. For each dish, TPEF intensity and FLIM images are taken under control conditions first from 3 different ROIs. We induce hypoxia (oxygen percentage is maintained below 1% as measured by the oxygen sensor) for 20 minutes then image 3 ROIs of the same dish with intensity and FLIM acquisitions. This experiment is performed using a 25x water immersion objective (N.A. 0.95). The results highlight that the NAD(P)H TPEF intensity contributions derived following the spectral crosstalk elimination approach exhibits the anticipated increase, resulting in a redox ratio decrease. Mitochondrial fragmentation is also increased under hypoxic conditions (Fig. S10D-H). Analysis of the g value distributions of the FLIM phasors indicates that hypoxia leads to significant decrease in the relative prevalence of mid G values and an increase in high G values, while there is a small but not significant decrease in the low G values. These are highly consistent with our attribution of changes in mid G values mostly to bound NADH, changes in high G values, to free NAD(P)H, and changes in low G values to bound NADPH (Fig. S10K-M).

**Supplementary Appendix**

**Supplementary Figures**

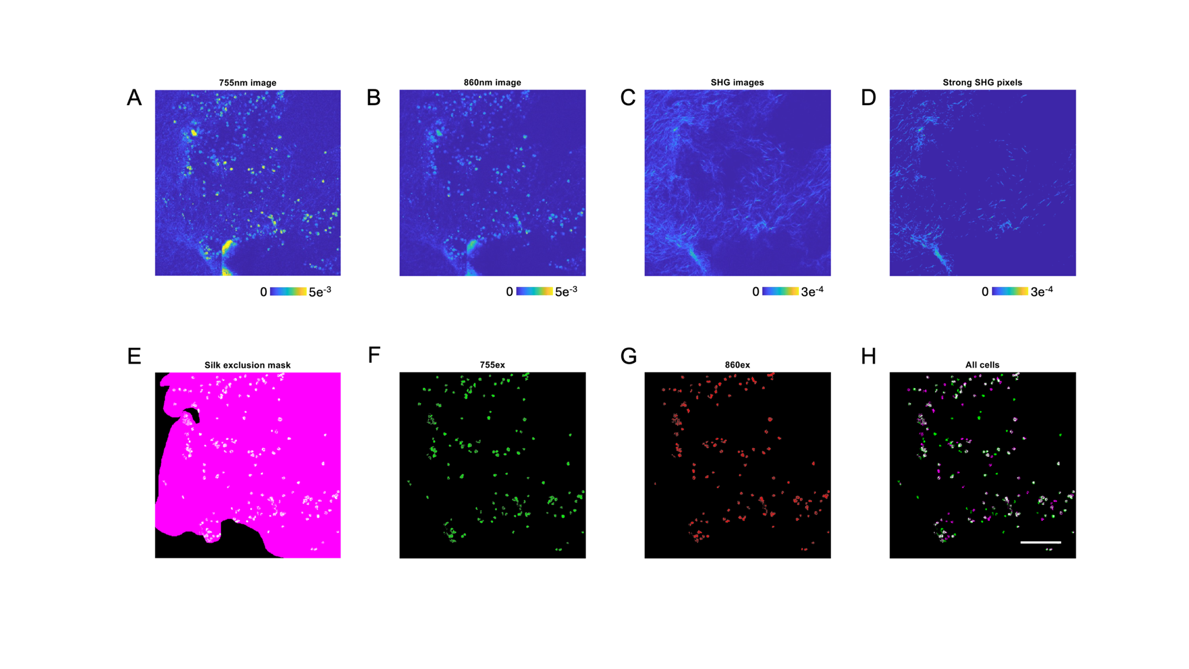

Fig. S1.

Workflow of cell feature extraction from spectral data. Integrated spectral image (2D) at (A) 755nm excitation, and (B) 860nm excitation from a neuronal monoculture 14 weeks in culture. (C) SHG image acquired at the same location at 860ex/430em. (D) Strong SHG signal with intensity larger than the 2^nd^ level of a three-level Otsu’s threshold extracted from (C). (E) Manually outlined silk exclusion map for this ROI, with the black region representing the excluded pixels. (F) Cell features identified from (A) using ISBA. (G) Cell features identified from (B) using ISBA. (H) Cell features identified from both wavelengths. Scale bar = 100μm.

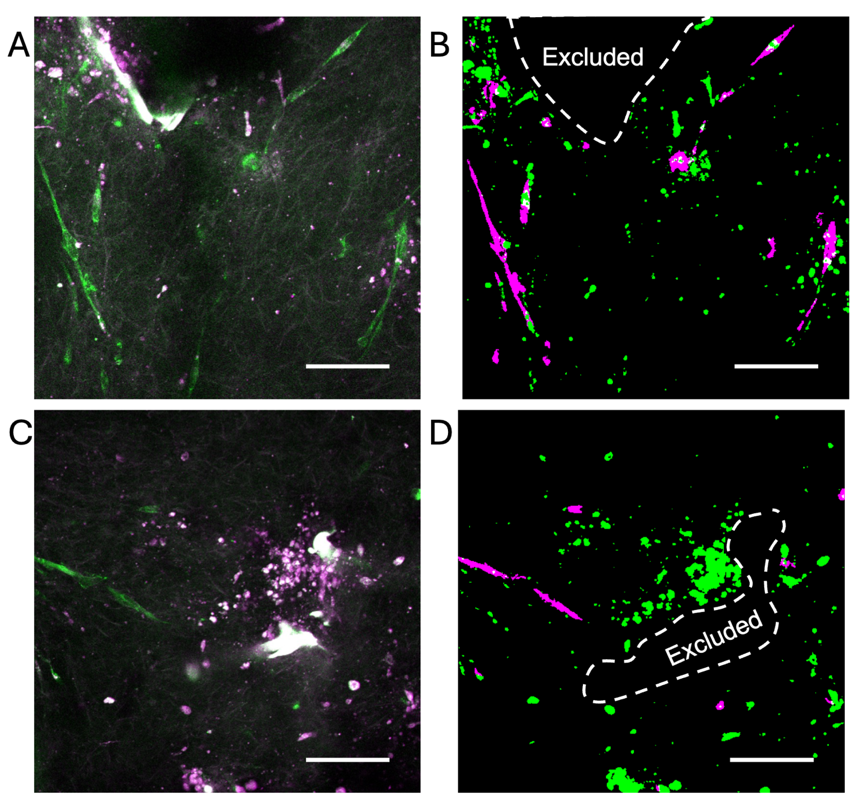

Fig. S2.

**ISBA based identification of neurons and glial cells in non-descanned images.** (A, B) Superimposed images acquired at 755EX/460EM (green) and 860EX/525EM (magenta) from two representative ROIs (C, D) Binary masks identified using ISBA for neurons (green) and glial cells (magenta). Since the images are maximum projections from 4 optical sections, if there are pixels from different cell features at the same x-y locations in different depths, the pixels are in white, showing overlap. Scale bar = 100μm.

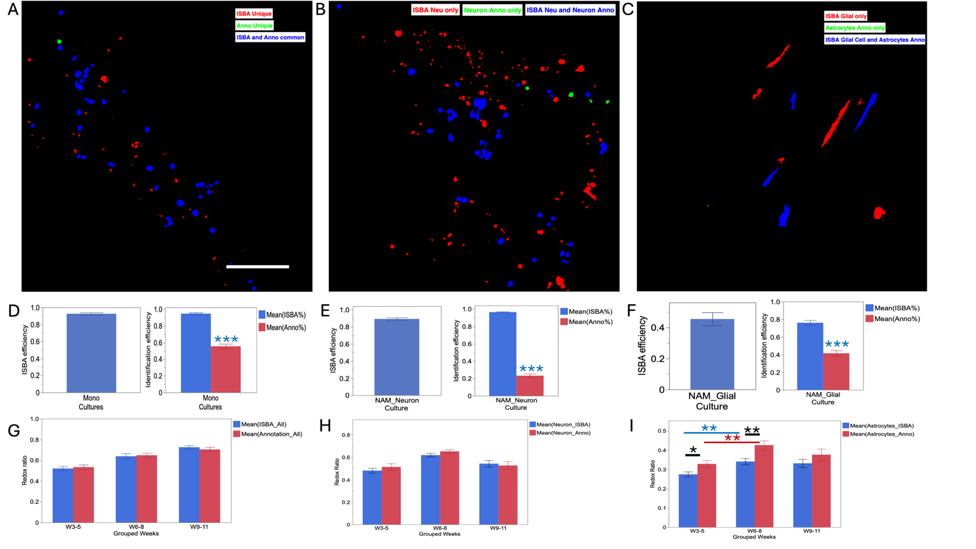

Fig. S3.

**Comparisons between ISBA and annotated objects.** Maximum projection of unique objects identified by ISBA (Red), manual annotations (Green), and common objects (Blue) in Monoculture data (A), Triculture neurons (B), and Triculture Glial cells (C). (D) ISBA efficiency for neuronal object detection in monoculture data, defined as the proportion of annotated neurons identified by ISBA. Identification efficiency represents the proportion of objects identified by each segmentation approach relative to the number of all objects identified by ISBA and annotations. (E, F) ISBA efficiency and identification efficiency for Triculture neurons (E), and for Triculture astrocytes (F). (G, H, I) Comparison of redox ratios calculated from all annotated monoculture neurons and ISBA objects (G), for triculture ISBA and annotated neurons (H), and triculture ISBA and annotated astrocytes (I). Note: Microglia are always identified using the mCherry label, so they are not included. * and ** represent p<0.05 and p<0.01 respectively. Scale bar = 100μm.

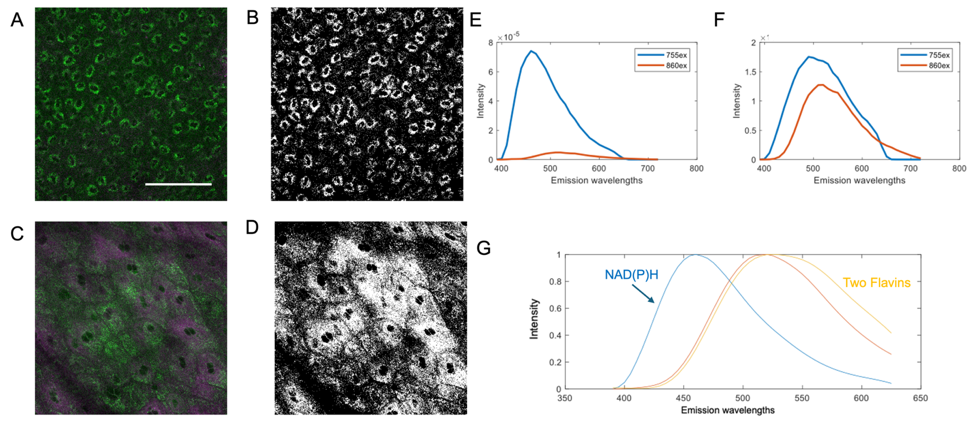

Fig. S4.

**Features for immature and mature mouse epithelial tissue.** A, C panels are overlays of the 755ex/460em and 860ex/525em images. (B, D) Isolated immature and mature epithelial regions used for spectral analyses. Mean spectra intensity from immature (E) and mature (F) epithelia at 755 nm and 860 nm excitation from the isolated cell regions. There is over 20 times stronger signal at 755nm than 860nm excitation in immature epithelia, making it a reliable NAD(P)H source, as NAD(P)H is barely detected at 860nm. (G) Spectrally deconvolved NAD(P)H and two additional components from analysis of epithelial spectra. Flavin shapes from epithelium tissues. Scale bar = 100 μm.

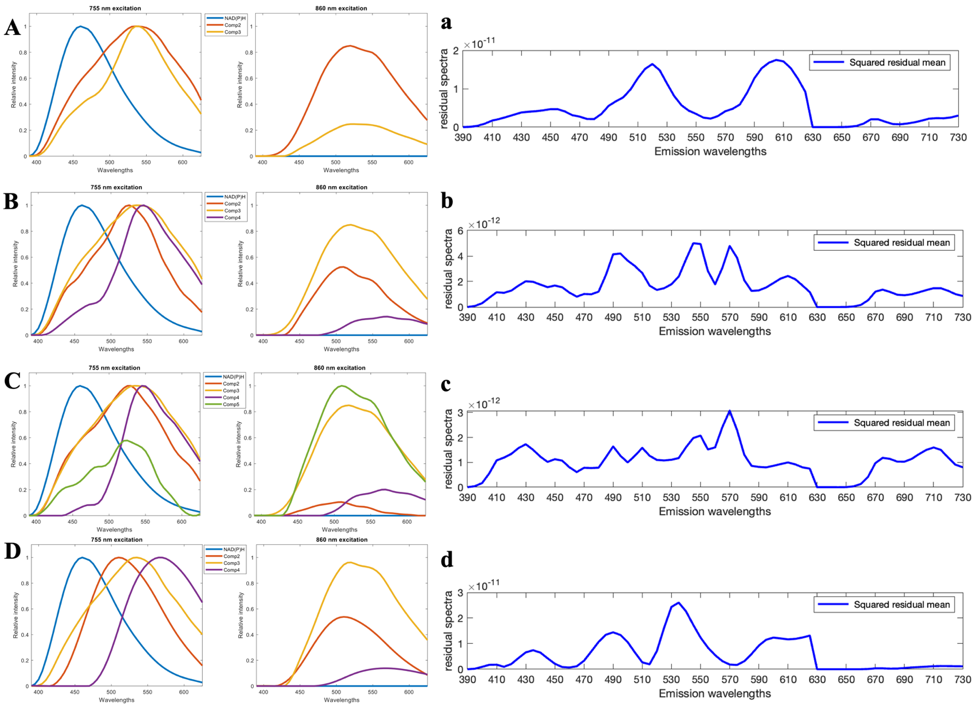

Fig. S5.

**Iterative spectral deconvolution approach for determining component spectra.** Iterative spectral deconvolution approach for determining component spectra. (A) Fixation of epithelial NAD(P)H in a three-component spectral deconvolution. (B) Four-component fit, yielding two fluorescent protein (FP) profiles and one lipofuscin profile at both 755 nm and 860 nm excitations. (C) Five-component fit. (D) Smoothed FP profiles used for 755 nm excitation in a subsequent four-component spectral deconvolution. Residual maps for the three-component (a), four-component (b), five-component (c), and four-component with modified FP profiles (d) fits. The three-component fit shows unresolved components in the residual maps. Starting from the four-component fit, the residual maps do not show additional shapes, indicating a good fit. We state this because the scale of the residual spectra is reduced from 10^^-11^ in the three-component fit to 10^^-12^ in the four-component fit.

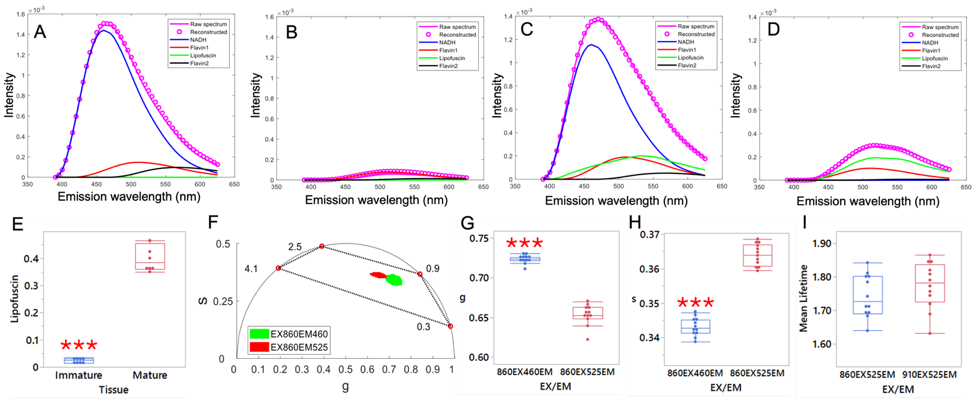
Fig. S6.

**Flavins in Epithelium tissues.** Fitting the epithelium data with brain-model-derived shapes provides good fits for both (A) immature and (B) mature epithelia. (C) Immature epithelia contain minimal lipofuscin as expected. (F) Phasors from immature epithelium at 860ex/460em and 860ex/525em. Reported lifetimes for LipDH (0.28 ns, 0.88 ns, and 4.14 ns) and FAD (2.47 ns) in tissue, are highlighted on the universal semi-circle. The extracted LipDH and FAD spectra indicate that the 460 nm detector captures mostly LipDH while the 525em detector captures LipDH and FAD information. The phasor shifts for the lifetime data collected from the 525 nm detector are consistent with more prevalent contributions from a 2.47 nm lifetime component, supporting enhanced sensitivity to FAD. These shifts are reproducible as indicated by smaller mean g value (G) and larger mean s values (H) from multiple immature epithelia FLIM images (pairwise two-tail t-tests). (I) The 910ex/525em FLIM phasors exhibit a trend towards higher long lifetime, consistent with an enhanced LipDH two-photon cross-section at 910 vs, 860 nm as reported by others.

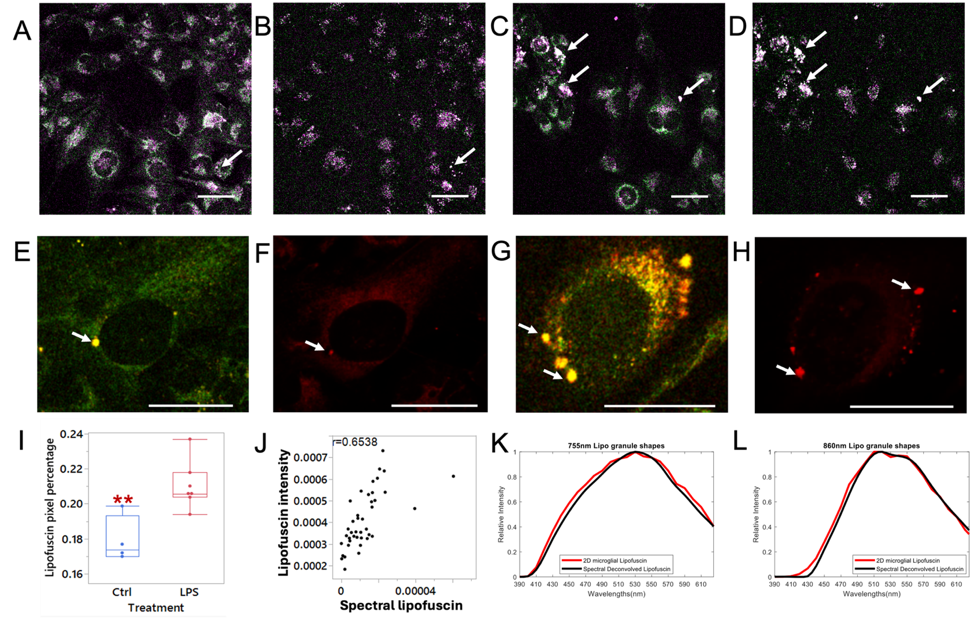

**Fig. S7. Lipofuscin Spectral Emission Validation.**

Superimposed autofluorescence images of HMC3 cells under control conditions are shown at 755 nm excitation, 460 nm emission (Green) and 525 nm emission (Magenta) in (A), and at 860 nm excitation, 460 nm, and 525 nm emission in (B). HMC3 cells incubated with lipopolysaccharide (LPS) for 24 hours to induce oxidative stress are imaged at 755 nm excitation, 460 nm, and 525 nm emission (C), and at 860 nm excitation, 460 nm, and 525 nm emission (D). Pixels with green color indicate stronger intensity at 460 nm emission, magenta indicates stronger intensity at 525 nm emission, and white pixels represent similar intensity at both emission channels, as highlighted by arrows pointing to lipofuscin granules that strongly fluoresce at all emission channels. Overlapped autofluorescence images at 755 nm excitation, 460 nm, and 525 nm emission of HMC3 cells from the control group after 24 hours of incubation before staining are shown in (E), and a confocal image after Nile blue A staining is shown in (F). Overlapped autofluorescence images of HMC3 cells from the LPS group after 24 hours of incubation are shown in (G), and after Nile blue A staining in (H). (I) shows the lipofuscin pixel coverage versus the cells, and (J) depicts the correlation of lipofuscin intensity and spectral lipofuscin amount in neuronal monocultures. Spectral data (2D, using ISBA) and intensity data (3D, using annotations) are from the same ROI from the same scaffolds, showing a significant correlation between spectral and intensity lipofuscin amount (p < 0.0001). Lipofuscin emission shapes from the 2D HMC3 culture and spectrally deconvolved shapes at 755 nm excitation (K) and 860 nm excitation (L) are also illustrated. Arrows in each image highlight the presence of lipofuscin granules. Asterisks *, ** indicate significant differences (p < 0.05, 0.01, respectively), with pairwise t-tests performed to assess significant differences between LPS and the corresponding control, as indicated by black asterisks. Scale bars in all panels represent 50 μm.

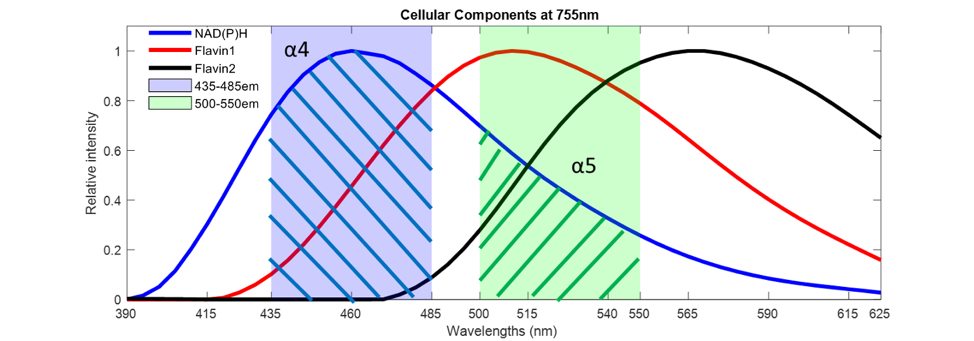

Fig. S8.

**Example of the crosstalk in the NAD(P)H channel.**

α4 is the AUC of NAD(P)H at 460em, shown by the region filled with blue lines, and α5 is the AUC of NAD(P)H at 525em, shown by the region filled with green lines. The same concept applies to other components.

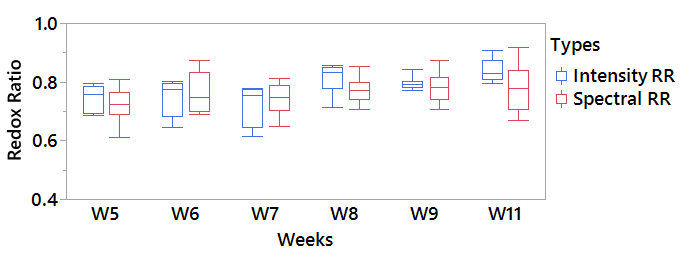

Fig. S9.

**Spectral redox ratio (2D, ISBA) and intensity redox ratio (3D annotations) comparisons.**

No significance between intensity RR and spectral RR is detected in any week.

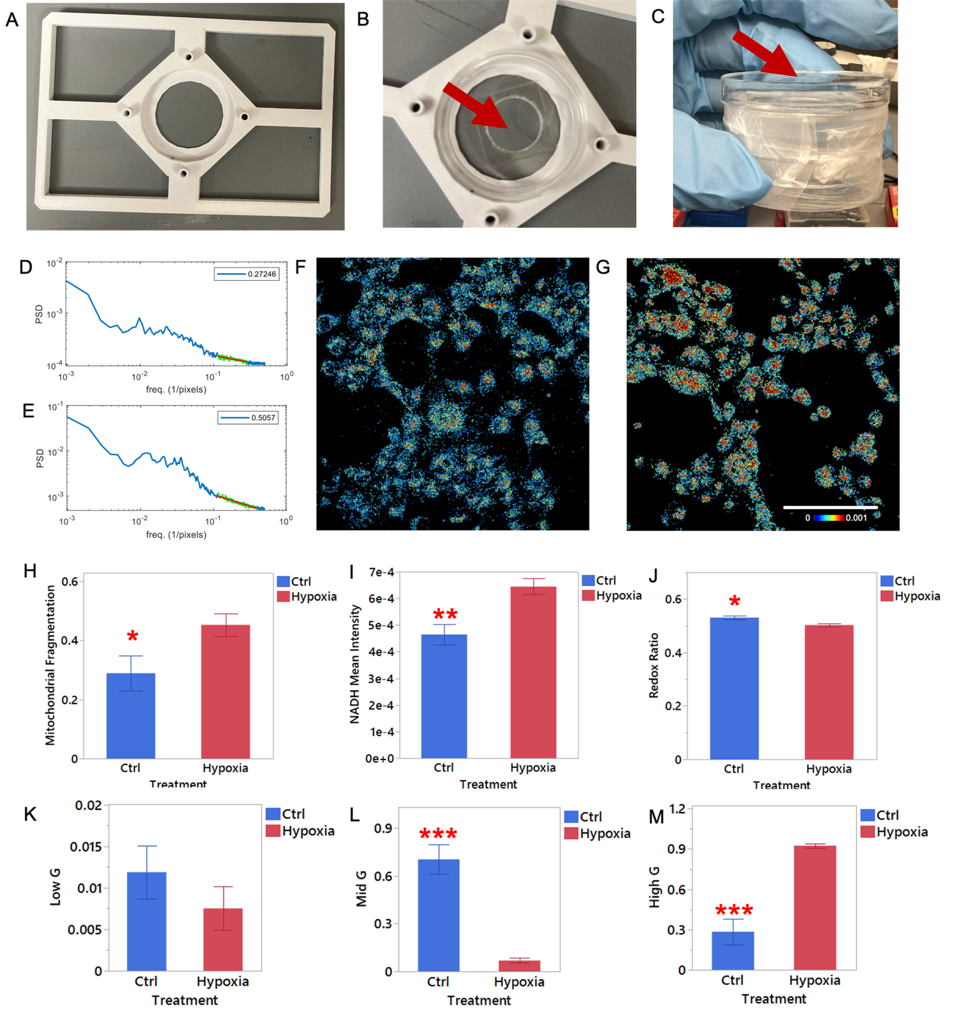

Fig. S10.

**Hypoxia experiment.**

(A) 3D-printed dish holder. (B) Top cap made by removing the glass bottom, allowing the placement of oxygen sensor, temperature sensor, and nitrogen tip as pointed by red arrow. (C) Cap placed on top of the glass-bottom dish with the sample, fully sealed. Mitochondrial fragmentation under control (D) and hypoxia (E) conditions. NAD(P)H maps under control (F) and hypoxia (G) conditions. (H) Mitochondrial fragmentation. (I) NAD(P)H intensity, normalized to the whole cell pixels per ROI. (J) Redox ratio based on the crosstalk-adjusted NAD(P)H and flavins amount. (K) Low G component. (L) Mid G component. (M) High G component. Panels F-G with the same color scale and scale bar = 100 μm. Pairwise two-tailed t-test is applied to H-M, with *, **, and *** representing p<0.05, p<0.01, and p<0.001 respectively.

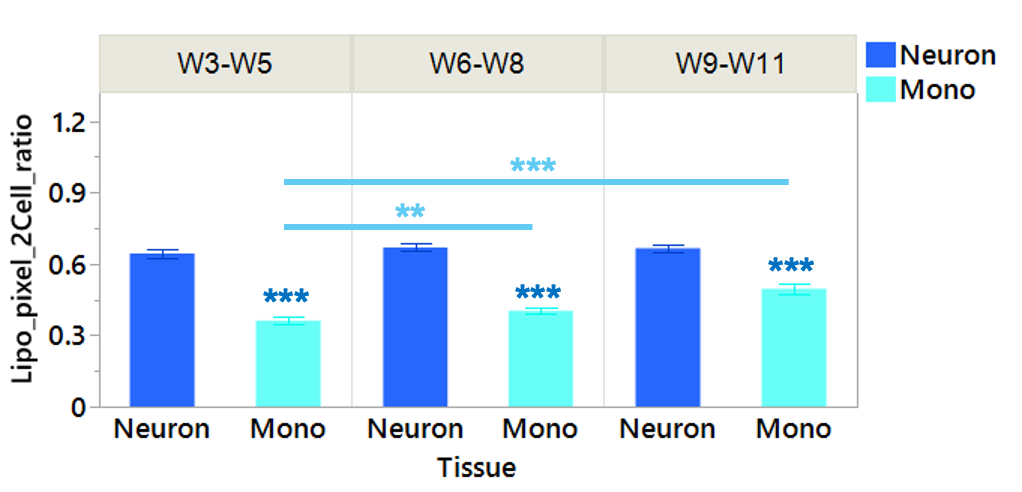
 **Fig. S11.**

**Lipofuscin versus cell pixel ratios of neurons in tricultures and monoculture neurons.**

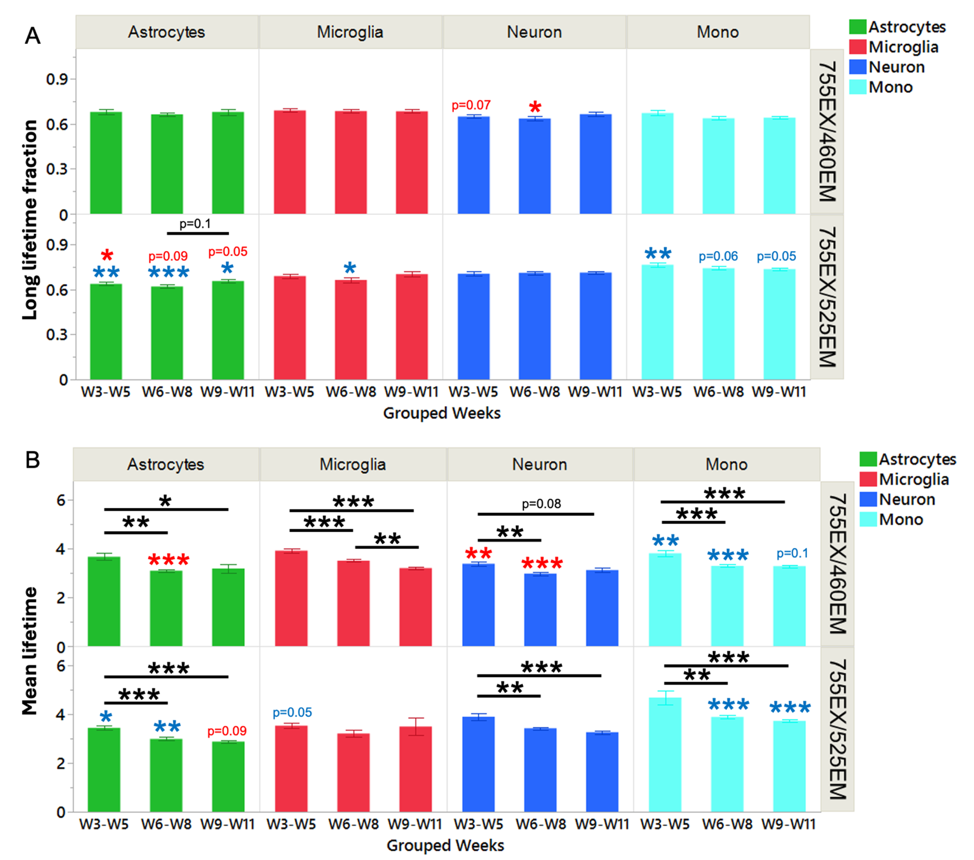

Fig. S12.

**Lifetime signature of distinct populations as a function of time.**

(A) Long lifetime fraction and (B) mean lifetimes for different cell features from the same ROIs at different emission channels. Each ROI is a data point. Bar graphs show mean values with standard deviation as error bars. Significance: *p<0.05, **p<0.01, ***p<0.001. Data is grouped into three ranges: Weeks 3-5, Weeks 6-8, and Weeks 9-11, with each cell type in a distinct color. Black asterisks indicate ANOVA results for each group across time points; colored asterisks show comparisons among neurons, astrocytes, and microglia within each group (ANOVA) and between neurons in monoculture and NAM (two-tailed t-test). The long lifetime fraction is calculated using linear line fitting to the phasor to identify cross-sections with the semicircle for long (left intersection) and short lifetime (right intersection) values. Mean lifetime is weighted by the amount of long and short lifetimes.

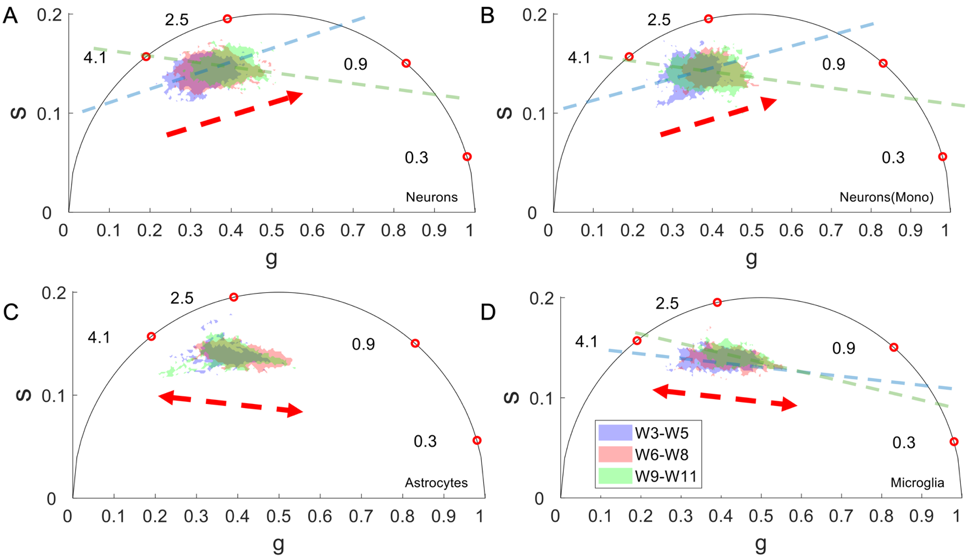

Fig. S13.

**Time-dependent phasor shifts in different cell types.** The phasor locations, integrate from all cell phasors acquired within Weeks 3-5 (blue shades), Weeks 6-8 (red shades), and Weeks 9-11 (green shades) for (A) Neurons, (B) Monoculture Neurons, (C) Astrocytes, and (D) Microglia. Neuronal phasors are rounder than glial phasors, showing a different trajectory of phasor shifts as indicated by red arrows. The dashed lines, which match the colors of the shades, represent linear fitting lines, highlighting the differences in long and short lifetimes (the intersection location with the semi-circle) between the early and later weeks. The red circles are reference lifetime locations.

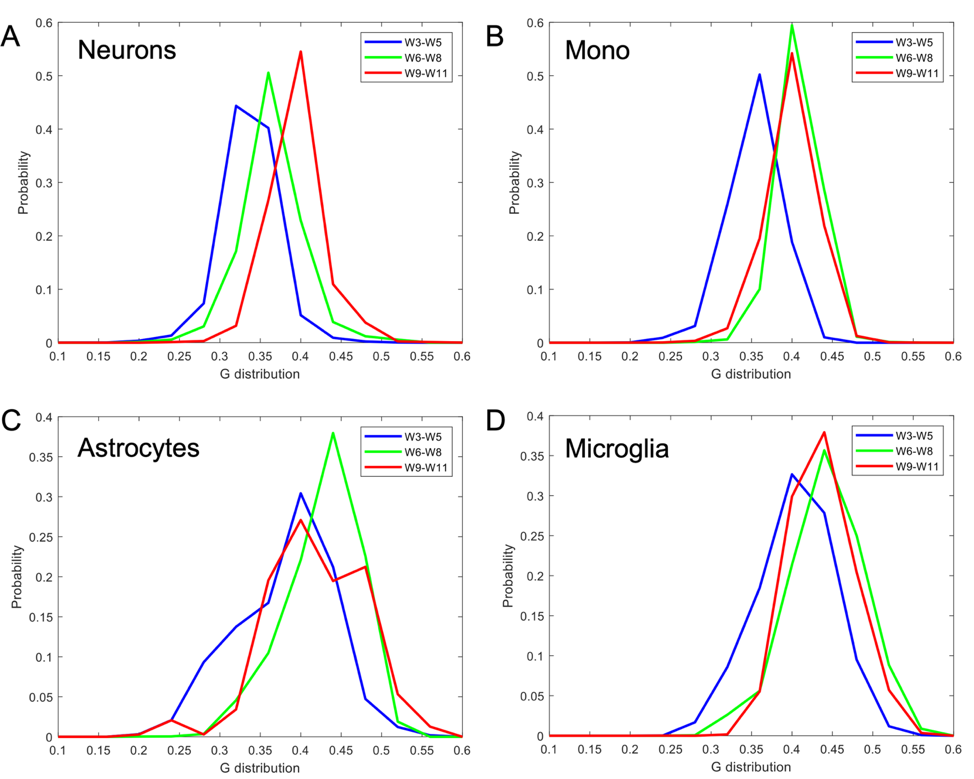

Fig. S14.

**Integrated g distributions.** Each colored line represents the integrated g distribution, created by transforming all g values from each phasor plots shown in Fig. 13 into a single distribution. These distributions summarize data from all ROIs acquired within Weeks 3-5 (blue), Weeks 6-8 (green), and Weeks 9-11 (red) for (A) Neurons, (B) Monoculture Neurons, (C) Astrocytes, and (D) Microglia. Each integrated g distribution is area under the curve (AUC) normalized to have a total probability of 1 for different g values. The superimposed distributions highlight a shift in g values towards larger g values, corresponding to a lower lifetime regime and a less bounded NAD(P)H pool. Glial cells show bi-directional shifts.

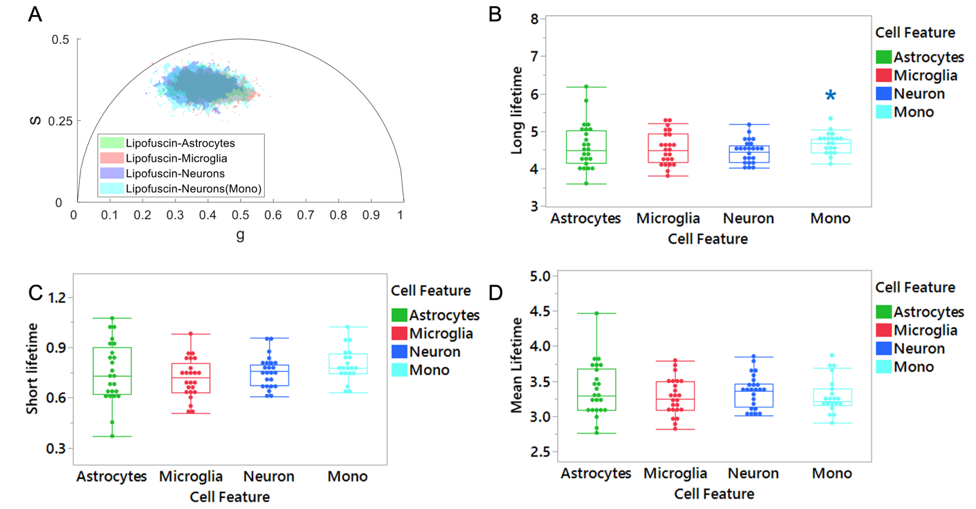

Fig. S15.

**Lifetime signatures of lipofuscin pixels from distinct populations.**

(A) Phasor locations integrated from all cell phasors across all weeks for each cell type at the 755ex/460em channel, with each cell type represented by a distinct color. (B) Long lifetimes, (C) short lifetimes, and (D) mean lifetimes are similar among astrocytes, microglia, and neurons in NAM. Only neurons in monoculture exhibit a longer long lifetime compared to neurons in NAM. Each data point includes all ROIs for the same scaffolds at each week to ensure sufficient lipofuscin pixels. Colored asterisks indicate comparisons among neurons, astrocytes, and microglia within each group (ANOVA) and between neurons in monoculture and NAM (two-tailed t-test). No significance was detected for C and D.

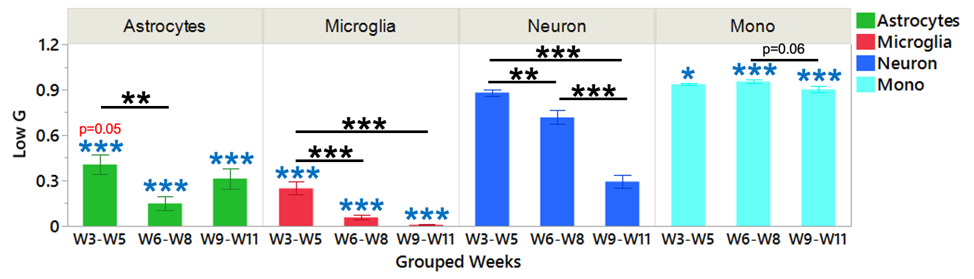
Fig. S16.

**NAD(P)H crosstalk channel with flavin information.**

755EX/525EM channel captures more flavins’ information relative to the 755EX/460EM channel. Free flavins contributes to higher Low G concentration, which is corresponding to lower G values in the phasor space.

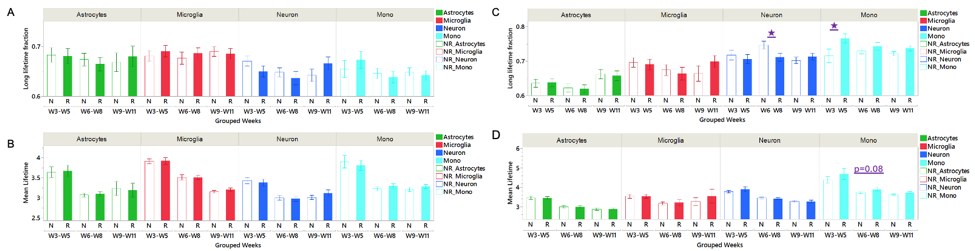
**Fig. S17.**

**Lipofuscin impact on FLIM readouts.**

A. Long lifetime fraction of each cell at 755ex/460em channel before and after lipofuscin removal. B. Mean lifetime of each cell at 755ex/460em channel before (NR: not removed) and after (R: removed) lipofuscin removal. C. Long lifetime fraction of each cell at 755ex/525em channel before and after lipofuscin removal. D. Mean lifetime of each cell at 755ex/525em channel before and after lipofuscin removal. Purple significance/stars represent the pairwise t-test before and after lipofuscin removal for each group. Other comparisons are not shown.

**Supplemental References**

1.	Liaudanskaya, V., et al., *Mitochondria dysregulation contributes to secondary neurodegeneration progression post-contusion injury in human 3D in vitro triculture brain tissue model.* Cell Death Dis, 2023. **14**(8): p. 496.

2.	Zhang, Y., et al., *Factors associated with obesity alter matrix remodeling in breast cancer tissues.* J Biomed Opt, 2020. **25**(1): p. 1-14.

3.	Varone, A., et al., *Endogenous two-photon fluorescence imaging elucidates metabolic changes related to enhanced glycolysis and glutamine consumption in precancerous epithelial tissues.* Cancer Res, 2014. **74**(11): p. 3067-75.

4.	Huang, C.X., et al., *Positron emission tomography imaging for the assessment of mild traumatic brain injury and chronic traumatic encephalopathy: recent advances in radiotracers.* Neural Regen Res, 2022. **17**(1): p. 74-81.

5.	Sanchez-Hernandez, A., C.M. Polleys, and I. Georgakoudi, *Formalin fixation and paraffin embedding interfere with the preservation of optical metabolic assessments based on endogenous NAD(P)H and FAD two-photon excited fluorescence.* Biomed Opt Express, 2023. **14**(10): p. 5238-5253.

6.	Baugh, L.M., et al., *Non-destructive two-photon excited fluorescence imaging identifies early nodules in calcific aortic-valve disease.* Nat Biomed Eng, 2017. **1**(11): p. 914-924.

7.	Pouli, D., et al., *Imaging mitochondrial dynamics in human skin reveals depth-dependent hypoxia and malignant potential for diagnosis.* Sci Transl Med, 2016. **8**(367): p. 367ra169.

8.	Liu, Z., et al., *Mapping metabolic changes by noninvasive, multiparametric, high-resolution imaging using endogenous contrast.* Sci Adv, 2018. **4**(3): p. eaap9302.
